# MyoRep: a novel reporter system to detect early muscle atrophy *in vitro* and *in vivo*

**DOI:** 10.1101/2025.02.27.640021

**Authors:** Andrea David Re Cecconi, Nicoletta Rizzi, Mara Barone, Federica Palo, Martina Lunardi, Mara Forti, Adriana Maggi, Paolo Ciana, Giulia Terribile, Michela Chiappa, Lorena Zentilin, Rosanna Piccirillo

## Abstract

Muscle atrophy occurs during physiological (i.e., fasting, aging) and pathological (i.e., amyotrophic lateral sclerosis, cancer) conditions and anticipates death. Since not all patients will undergo muscle wasting, it would be highly useful to identify them soon to intervene early. We have studied the promoters/enhancers of a subset of atrophy-related genes or atrogenes upregulated in muscles of rodents during various kinds of atrophy (i.e., disuse, diabetes, cancer, fasting, uremia) for their ability to induce early atrophy. Comparing their upstream non-coding regions, using as backbone *MuRF1* promoter (one of the earliest muscle-specific genes induced by wasting), we cloned various promoters upstream of a doubled reporter system (FLuciferase/tdTomato). Through *in vitro* and *in vivo* studies in mice subjected to denervation or cancer injection, we selected a sequence (i.e., MyoRep) able to predict atrophy upon cut of the sciatic nerve or cancer. *In vivo* imaging of MyoRep mice emit a bioluminescent signal earlier than muscle loss. Importantly, MyoRep was unable to sense atrophy during fasting or physiological variations following the circadian rhythms. Since MyoRep can discriminate muscle loss due to pathological conditions from physiological ones anticipating wasting, it represents an unprecedented tool to predict it early in various diseases with local or systemic atrophy.

## INTRODUCTION

Skeletal muscle atrophy consists of progressive loss of muscle proteins due to an imbalance between protein generation and degradation. Atrophy occurs physiologically under certain circumstances as upon bed rest or fasting or following circadian rhythms. To a major extent, patients can face progressive muscle wasting during local or systemic diseases, such as loss of innervation or cancer, respectively. Other diseases causing severe muscle wasting are spinal cord injury with subsequent paralysis and disuse atrophy, Amyotrophic Lateral Sclerosis, burn injury, heart or kidney failure, sepsis and muscular dystrophies.

Muscle wasting is one of the hallmarks of cachexia (a debilitating condition involving involution of multiple tissues). Moreover, it is a major medical need since it reduces the response to therapies and leads to premature death that usually arises when lean muscle mass loss reaches 30-40% of body weight ^1^. The overall prevalence of cachexia is quite high and increasing in industrialized countries. It is estimated that cachexia affects about 9 million patients, which is 1% of all patients with any disease ^2^.

Paradoxically, despite skeletal muscle is among the most abundant tissues in our body, it is one of the most “undrugged” tissues, for which less drugs exist to repair or spare it. Either during physiological and pathological atrophies, the expression of a common set of genes, namely atrogenes, is changed in muscles ^3^. Among those that are early upregulated in various types of atrophy, there are genes encoding for transcription factors as FoXO3 or muscle-restricted ubiquitin ligases as atrogin-1 and MuRF1 ^4,5,6^ that promote muscle protein degradation through the proteasome.

In the attempt to generate a reporter mouse able to emit a bioluminescent signal under a muscle-specific promoter to follow *in vivo* only pathological atrophy, over the years we have analysed either binding sites for specific transcription factors driving atrophy (as FoXO3, NF-kβ ^7^, STAT3 ^8^, Smad 2/3 ^9^) or promoters of atrogin-1 or MuRF1 in *in vitro* luciferase-based assays. Even if other reporter mice have been generated in the past as able to sense and signal muscle wasting ^10,11^, we believe that ours, namely MyoRep (i.e., Myo standing for muscles and Rep for reporter) based on a much more improved version of the MuRF1 promoter is better for two main reasons. Firstly, MyoRep is able to sense only pathological atrophy (caused by cut of the sciatic nerve or cancer) and not physiological one (as that induced by food deprivation), because such promoter has been deprived of GRE (glucocorticoids-responsive elements). Secondly, its precocity in sensing atrophy has been obtained also by repeating in tandem the binding sites for specific transcription factors (as TWIST ^12^, FoXO3 and myogenin-binding sites ^13^), highly reinforcing the responsiveness of such newly engineered promoter.

To assess the robustness of MyoRep promoter, we have evaluated its activity in *in vitro* luciferase assays, demonstrating its ability to drive the expression of a reporter gene (Firefly Luciferase) prior to protein loss, so before myoblast atrophy. Additionally, MyoRep functionality has been confirmed in *in vivo* experiments. By transfecting such reporter plasmids in muscles of adult mice, or by locally injecting AAV9 vectors carrying the MyoRep sequence upstream of Firefly Luciferase in leg muscles, we demonstrate its early activation and its ability to longitudinally track over time muscle wasting in mice by *in vivo* imaging. The generated MyoRep mouse expressing the cassette only in skeletal muscles and heart is able to emit a local bioluminescent signal easily detectable by *in vivo* imaging upon muscle denervation or sarcoma MCG101 injection. This is in line with the 3R principle by minimizing animal use, reducing experimental variability and refining methodologies to obtain more physiologically relevant data through non-invasive *in vivo* imaging ^14^.

Overall, we believe MyoRep mouse can be a useful tool to identify novel drugs against atrophy and/or useful early biomarkers or to better understand the dynamics behind muscle wasting and which muscles among others are preferentially lost during various disease without the need of sacrificing many mice to weight their muscles.

## MATERIAL AND METHODS

### Cell lines

C2C12 (ATCC, Manassas, VA, USA) is an immortalized mouse myoblast cell line obtained from the C3H mouse strain. It was grown in DMEM (Dulbecco’s Modified Eagle’s Medium, Gibco, Waltham, MA, USA), supplemented with fetal bovine serum (FBS) (Euroclone, Pero, Italy) and 2 mM L-glutamine, and maintained in culture at 37°C with 5% CO_2_. C26 is a colorectal adenocarcinoma cell line obtained from BALB/c mice, grown in DMEM (Dulbecco’s Modified Eagle’s Medium, Gibco, Waltham, MA, USA), supplemented with 10% FBS and 2 mM L-glutamine at 37 °C with 5% CO_2_. C26 cells were kindly gifted by Prof. Mario Paolo Colombo (IRCCS-Istituto Nazionale dei Tumori, Milan, Italy). MCG101 is a sarcoma cell line, grown in McCoy’s 5A (Gibco) medium with 10% FBS and 2 mM L-glutamine at 37°C with 5% CO_2_. MCG101 cells were shared by Prof. Anders Blomqvist (Linköping University, Linköping, Sweden). 4T1 cells were grown in Ham’s F12 (Gibco) with 10% FBS and 2 mM L-glutamine, at 37°C with 5% CO_2_. All cell lines were not contaminated by mycoplasma.

### Plasmids

MuRF1 promoter was cloned into the vector Pgl4.10[luc2] (Promega, Madison, WI) upstream of the *Firefly Luciferase* gene, with the cloning sites XhoI/EcoRV (Genescript, Piscataway, New Jersey, USA).

We used the following plasmids, all engineered by us:

- m(urine)MuRF1_pGL4.10[luc2], simply referred to as mMuRF1;
- mMuRF1-TWIST_pGL4.10[luc2], simply referred to as TWIST;
- mMuRF1-GREDEL_pGL4.10[luc2], simply referred to as GREDEL.

All the plasmids were co-transfected with pRL-TK-Renilla, a plasmid encoding for *Renilla Luciferase*, used as index of transfection efficiency to normalize the data (Promega, Madison, WI, USA) with a ratio of 1:50 (Renilla:Firefly).

The following plasmids are referred to the Table I:

- pGL3-FHRE ΔXRE: it contains 3 FoXO responsive elements without xenobiotic responsive ones, received from Prof. Klotz, University of Alberta, Canada;
- PGL4.10 2D4F: it contains 4 FHRE and 2 DBE sequencing, engineered by us;
- pGL3-DBE (Daf-16 binding elements): it contains six Daf-16 binding elements, the ortholog of FOXO factors in *C. elegans*, received from Prof. M. Sandri, University of Padova, Italy;
- pGL3-pAT1 3.5 Kb (Atrogin-1 promoter), received from Prof. A.L. Goldberg, Harvard Medical School, Boston, USA;
- pGL3(CAGA)12 Firefly LUC (SMAD 2/3 binding sites): it contains 12 copies of (CAGA) box upstream of the minimal Adenovirus major late promoter (MLP) (TATA box + initiation sequence of MLP) controls the expression of FLuc, received from Prof. M. Sandri, University of Padova, Italy;
- pGL3 - 5 Kb MuRF1 promoter, received from Prof. A.L. Goldberg, Harvard Medical School, Boston, USA;
- 4XM67-Luc-STAT3: it contains 4 STAT3 (M67) binding sites, received from Prof. D. Bessel, Max-Delbrück-Centrum für Molekulare Medizin, Berlin, Germany;
- pNF-kβ Luc2-IRES-EGFP, as in ^15^;
- pGL4.30(Luc2P/NFAT-RE/Hygro), Promega, cat. n° E8481,
- TK-LXRE3-LUC, it contains 3 LXRE copies, received from Prof. A. Moschetta, Università “Aldo Moro”, Bari, Italy;
- pGL4.33(Luc2P/SRE/Hygro), Promega, cat. n° E1340;
- pGL4.29(Luc2P/CRE/Hygro), Promega, cat. n° E8471.

**Table I:**
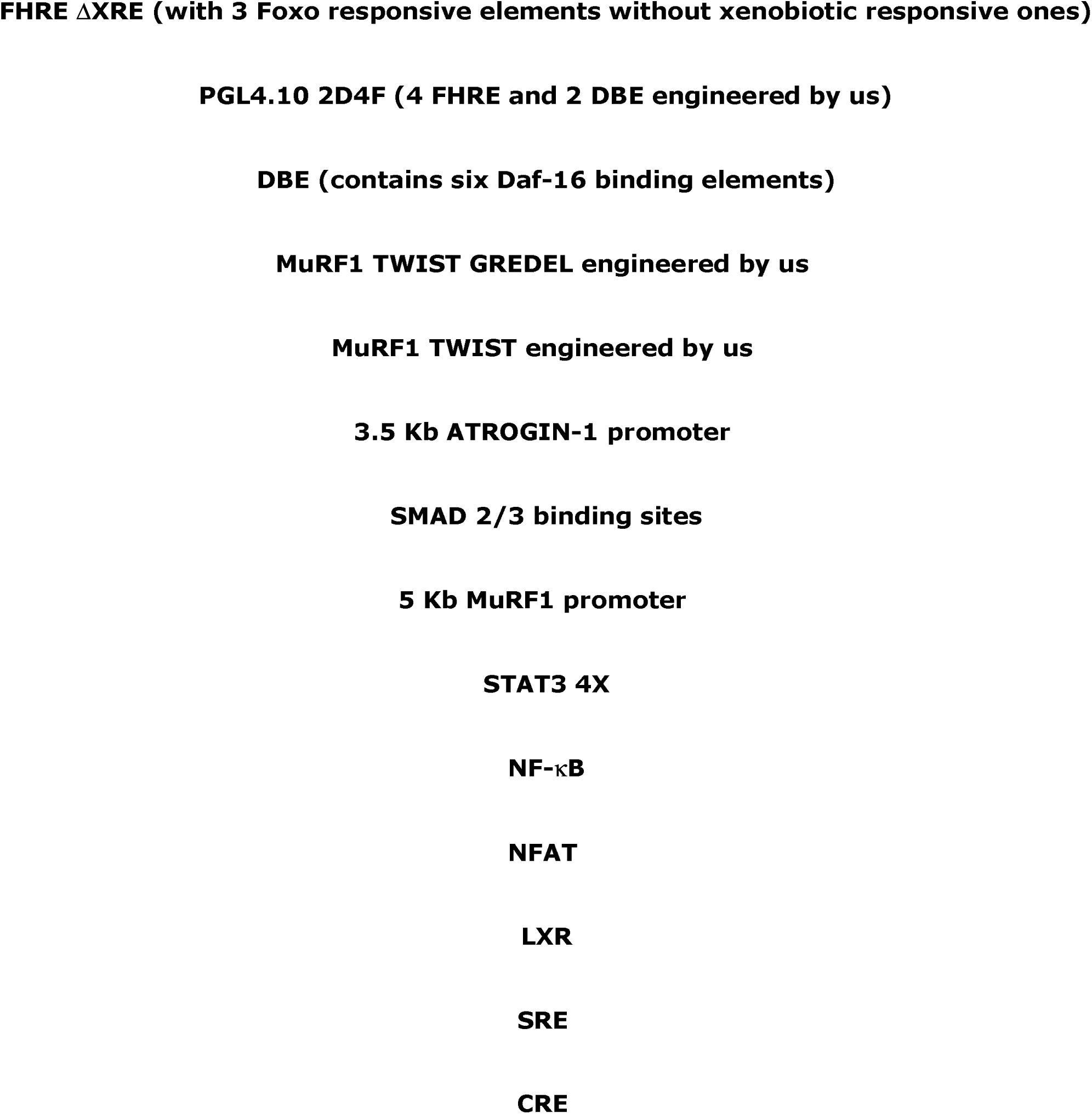
A list of the regulatory sequences tested *in vitro* in Lucifease-based assays is provided.

### Cell transfection

Twenty-four hours before transfection, C2C12 cells were seeded at 17,500 cell/cm^2^. We used 48 well-plates for Luciferase-based assays (see later in the section). C2C12 cells were transfected using Lipofectamine 2000 (Invitrogen, Waltham, MA, USA), according to manufacturer’s instructions.

### Cell treatments

C2C12 cells transfected with the above-mentioned plasmids were treated with conditioned medium of tumoral cells. Briefly, tumoral cells as C26 and 4T1, were grown in their medium. After 48 hours, conditioned media were harvested when cells were 80-90% confluent, centrifuged for 5 minutes at 1,000 rpm. Such supernatant was then used to treat C2C12 for 24 hours. Transfected C2C12 cells were treated with dexamethasone for 24 hours, at the concentration of 1 and 10 µM.

### *In vitro* and *ex vivo* Luciferase-based assays

Luciferase assay was done on C2C12 cells grown in a 48 well-plate and transfected using Lipofectamine 2000 with different combination of plasmids, in the following relative proportions:

- mMuRF1-TWIST – pRL-TK-Renilla ratio 50:1;
- mMuRF1-GREDEL– pRL-TK-Renilla ratio 50:1;
- mMuRF1– pRL-TK-Renilla ratio 50:1.

The next day, cells were treated with dexamethasone or media conditioned by tumoral cells. Firefly and Renilla luciferase activities were measured in cell lysates using the Dual-Luciferase Reporter Assay System (Promega, Madison, WI, USA) and the signal was quantitated using a luminometer (Glomax 20/20 single tube luminometer, Promega, Madison, WI, USA). Luciferase assay was also made *ex vivo*, on muscles excised from MyoRep mice injected with MCG101 tumor or C57BL/6J-Apc^Min/+^ injected with MyoRep-AAV9 in Tibialis Anterior (TA) for 10 weeks. Muscles were lysed with Passive Lysis Buffer (PLB) and firefly luciferase activity was measured using the Luciferase Assay kit (Promega). The signal was quantitated using the Glomax 20/20 single tube luminometer and related to total protein content of the sample measured with Bradford.

### *In vivo* experiments

*In vivo* experiments have been carried out on C57BL/6J-MyoRep mice and C57BL/6J-Apc^Min/+^ and *wild-type* littermates (The Jackson Laboratory - Charles River Italia, Calco, Italy). Four animals have been housed per cage in standard conditions with unlimited access to food and water, with 12 hours of light and 12 hours of dark. Inside the cage, environmental enrichment was provided. Males and females have been separated into different cages. Mice have been identified with a hole in different positions of the ears. Procedures involving animals and their care were conducted in conformity with institutional guidelines in compliance with national and international laws and policies. The Mario Negri Institute for Pharmacological Research IRCCS (IRFMN) adheres to the principles set out in the following laws, regulations, and policies governing the care and use of laboratory animals: Italian Governing Law (D.lgs 26/2014; Authorisation n° 19/2008 - A issued 6 March 2008 by Ministry of Health); Mario Negri Institutional Regulations and Policies providing internal authorisation for persons conducting animal experiments (Quality Management System Certificate - UNI EN ISO 9001:2015 - Reg. n° 6121); the National Institutes of Health (NIH) Guide for the Care and Use of Laboratory Animals (2011 edition) and European Union (EU) directives and guidelines (European Economic Community (EEC) Council Directive 2010/63/UE). The Statement of Compliance (Assurance) with the Public Health Service (PHS) Policy on Human Care and Use of Laboratory Animals has been recently reviewed and will expire in 2027 (Animal Welfare Assurance #972/2020PR - 157/2023PR - 992/2023PR).

### C57BL/6J-Apc^Min/+^ mouse model

In C57BL/6J-Apc^Min/+^ (Apc^Min/+^) mice, colorectal cancer-induced cachexia is evident around 12-14 weeks of age and their average lifespan is about 20-25 weeks. To genotype them, we proceed with a PCR using a little biopsy from the mouse ear, using the primers in the following table:

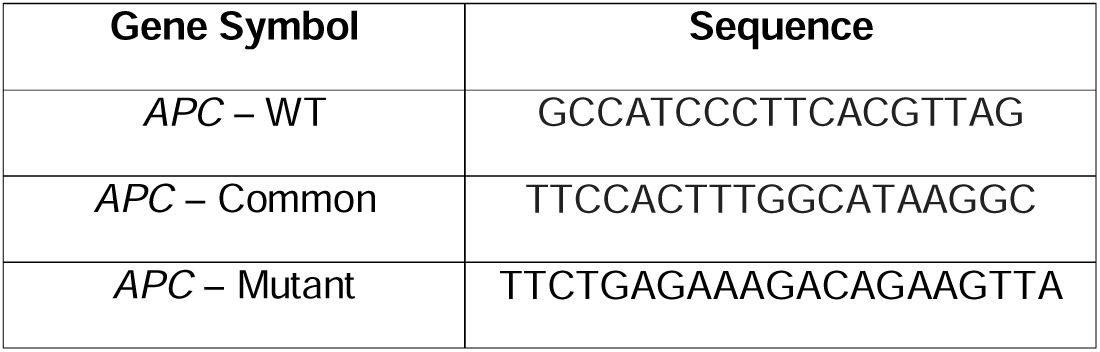

Apc^Min/+^ mice have been weighted once a week starting from 9-10 weeks of age, then, when the first signs of cachexia appear, later 3 times a week, until euthanasia.

### MCG101 and 4T1 mouse models

MCG101 cells were seeded at a density of 12,000 cells/cm^2^ and 4T1 cells at 25,000 cells/cm^2^ in a filtered-capped flask. Forty-eight hours later, cells were detached using trypsin-EDTA 0.25% (Gibco, Waltham, MA, USA) and counted through Beckman Coulter Counter. Then, they were centrifuged for 10 minutes and the pellet was resuspended in sterile PBS, to obtain 0,5 × 10^6^ cells in 200 µl of sterile PBS (phosphate buffered saline) for MCG101 and 2 × 10^5^ for 4T1 for each mouse. MCG101 cells were subcutaneously injected into the upper right flank of C57BL/6J or C57BL/6J-MyoRep male mice of 10-12 weeks of age using a 1 ml syringe with a 25 Gauge-needle. 4T1 cells were subcutaneously injected into the upper right flank of BALB/c male mice of 10-12 weeks of age (Envigo, Indianapolis, IN, USA) using a 1 ml syringe with a 25 Gauge-needle. The control groups received 200 μl of sterile PBS.

### MyoRep reporter mouse

In the MyoRep plasmid the promoter was cloned into the NF-Kβ-loxP-STOP1x-loxP-luc2-ires-tdTomato plasmid by substituting the NF-Kβ responsive promoter with MyoRep promoter in the XhoI/AscI restriction sites by using standard cloning procedures. In this way, we obtained pTargeting-MyoRep ^15^. In order to identify the best promoter sequences and responsive elements we analysed a panel of atrogenes paying attention to conserved responsive elements included in the majority of atrogenes and also the sequences of responsive elements typical of the early atrogenes. At the end of such bioinformatics analysis, the sequence identified as the best one to do MyoRep system was derived from MuRF1 promoter, but we deleted some regions and dimerized a short fragment that holds several well-conserved binding sites: E-Boxes (EM1, EM2 and EM3) binding sites for Twist; FOXO, Smad and myogenin binding sites as indicated in the sequences below:

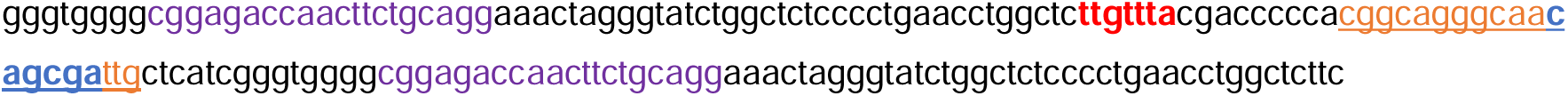
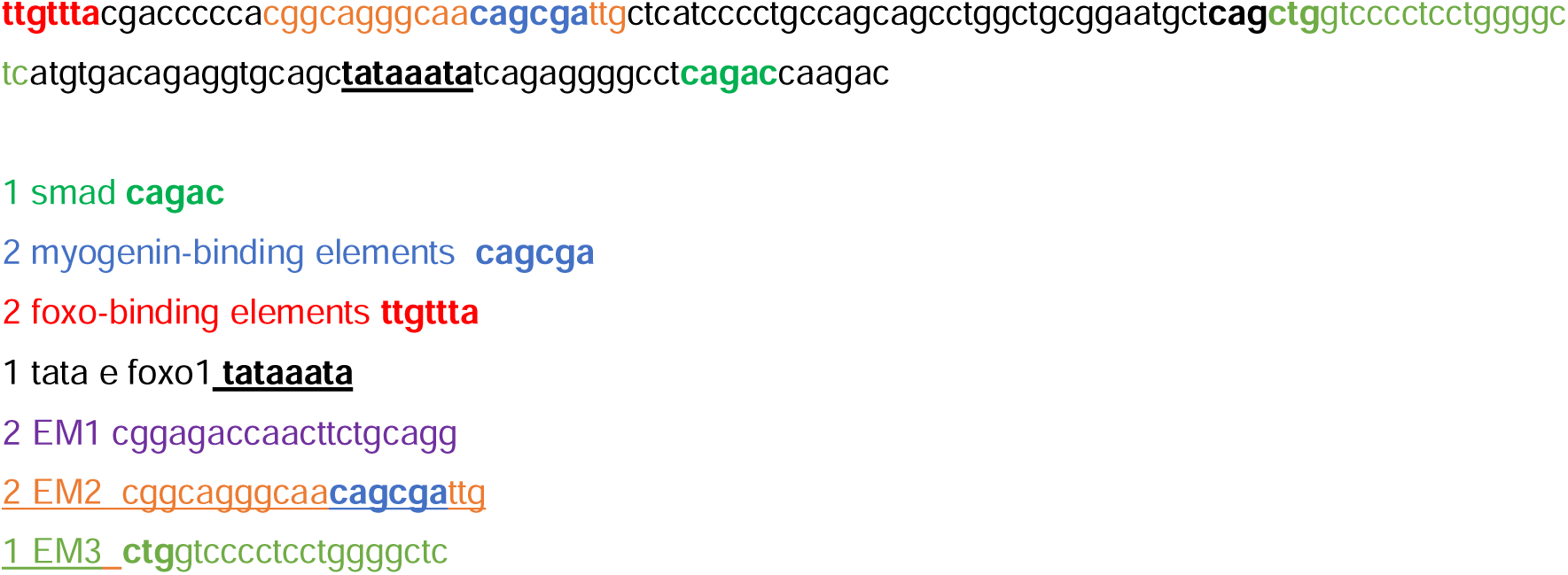

The targeting vector MyoRep was linearized with NotI and transferred into sv6.4 embryonic stem cells by electroporation: 35 µg/DNA each using 15 million cells (Core Facility for Conditional Mutagenesis, DIBIT San Raffaele, Milan, Italy). Positive clones were selected with puromycin (1 µg/μl). More than four hundred resistant clones for each transgene were screened for homologous recombination by PCR. One positive clone was injected into C57BL/6NCrL blastocysts which are transferred to pseudo-pregnant CD-1 females. We obtained 9 chimeric male mice (with 80–90% of chimerism) that were mated to wild-type (WT) C57BL/6J female mice to produce F1 MyoRep-stop transgenic mice. The stop sequence in MyoRep-stop (Luc2) mouse is removed using the loxP system after crossing with the B6.Cg-Tg(ACTA1-cre)79Jme/J mouse (The Jackson Laboratories, USA). Compared to the Luc2 mouse, the MyoRep mouse shows a basal bioluminescence signal following the removal of the stop sequence (**Supplementary Fig.3**).

To genotype CRE and Luc2 mice, we proceed with a PCR using a little biopsy from the mouse ear, using the primers in the following table:

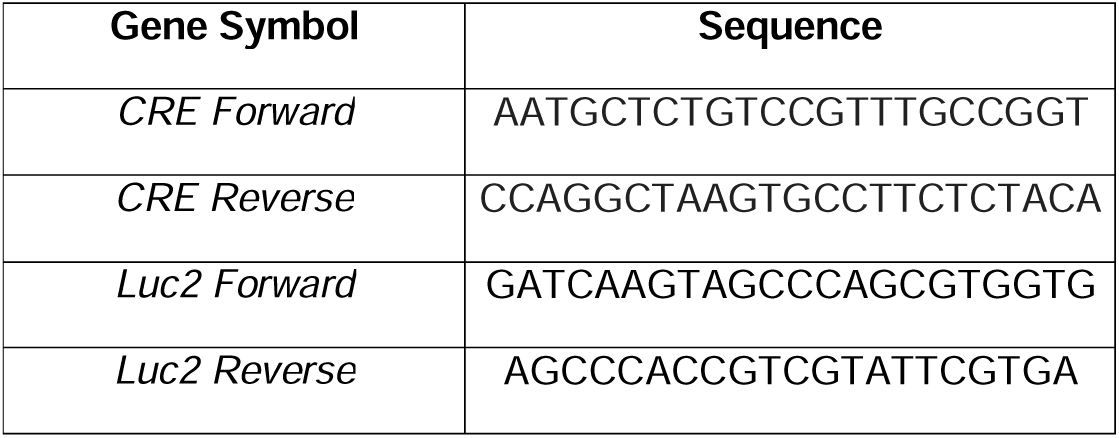

Instead, the sequence of the so-called mMuRF1-TWIST promoter is as follows:

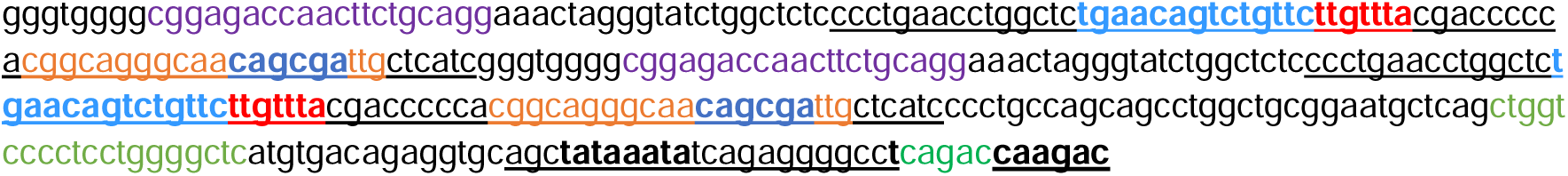
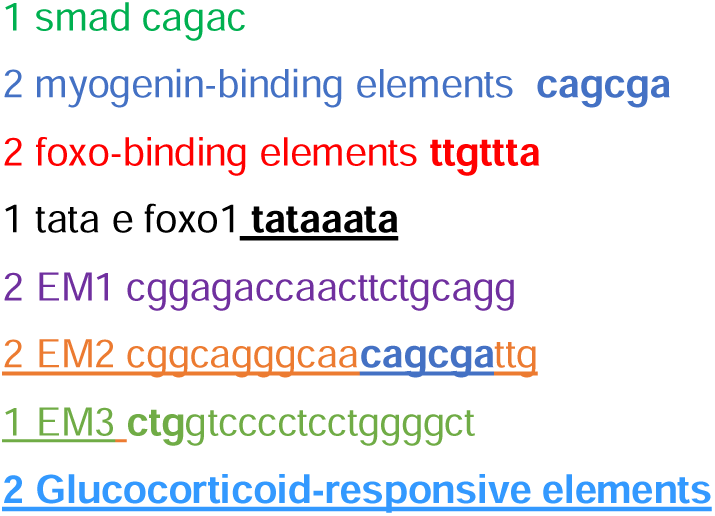

Finally, the sequence of the so-called mMuRF1 promoter is as follows:

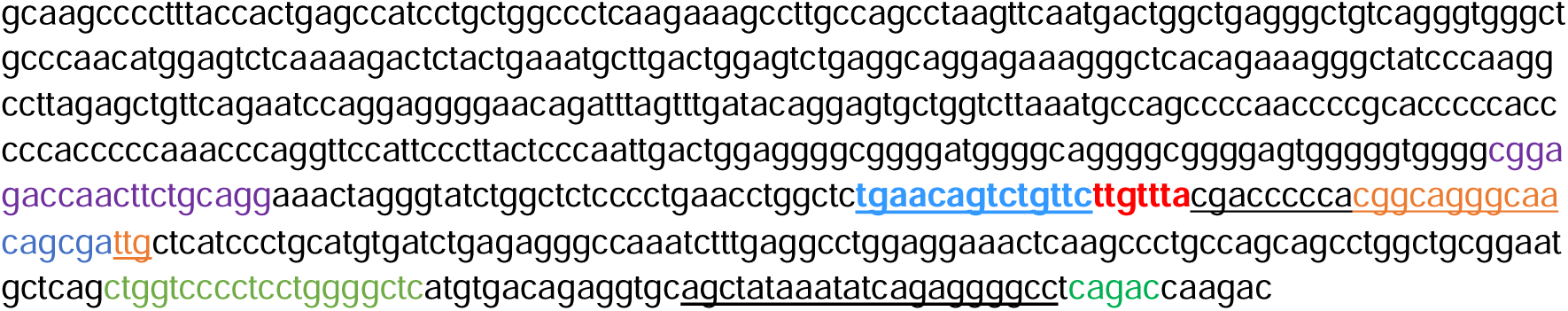

### Muscle electroporation with plasmids

The Endotoxin-free Maxi Prep kit (Invitrogen, Carlsbad, CA, USA) was used to purify plasmids (TWIST and GREDEL) from bacteria for muscle electroporation. The animals were anesthetized by inhalation of 3% isoflurane and 1% O_2_. Their hindlimbs were shaved and a linear incision of about 1 cm was made in the skin to expose the TA. One flatted electrode was then placed under the muscle, and using a 30-gauge Hamilton syringe, 20 µg of plasmid DNA in 30 μl of 0.9% NaCl solution in H_2_O were injected in the muscle. The second flatted electrode was then placed over the muscle, which was electroporated with five pulses (21 V) of 20 milliseconds (ms) each, with a 200 ms interval. Electroporation was done with the BTX ECM 830 Square Wave Electroporation System (Harvard Apparatus, Cambridge, MA, USA). Finally, the wound was sutured and disinfected with betadine.

### MyoRep-AAV9

Adeno-associated viruses 9 (AAV9) that express the MyoRep construct or musclin were injected intramuscularly into TA of Apc^Min/+^ mice or WT mice at 10 weeks of age of both sexes the. Based on our preliminary data not shown, we set out to use the concentration of 10^12 vg/ml of such AAV9 in 30 µl of PBS per leg of mouse. Recombinant AAV9 vectors used in this study were prepared by the AAV Vector Unit at the International Centre for Genetic Engineering and Biotechnology Trieste. Briefly, infectious AAV vector particles were generated in HEK293T cells cultured in roller bottles by a three-plasmids transfection cross-packaging approach whereby the vector genome was packaged into AAV capsid serotype-9 ^16^. Purification of viral particles was obtained by PEG precipitation and two subsequent CsCl_2_ gradient centrifugations ^17^. The physical titer of recombinant AAVs was determined by absolute quantification of vector genomes (vg) packaged into viral particles, by real-time PCR.

### In vivo imaging

WT mice after TWIST and GREDEL electroporation or AAV9 injection in TA and MyoRep mice emit a bioluminescent signal that can be easily detected through *in vivo* imaging (IVIS machine, Perkin Elmer, Milan, Italy). Averagely once a week, we performed an IVIS scan. We injected a volume of luciferin (Perkin-Elmer – 20 mg/kg) and 5 minutes later, we anesthetized mice in the induction chamber, shaved the hind legs, as the hair shields the bioluminescence, and we placed them into the instrument, in dorsal position, exposing the TA and gastrocnemius (GAS) muscles of both legs as best as possible, or in ventral position. Ten minutes after the luciferin injection, we performed one minute-scan for bioluminescence (for plasmids electroporation and AAV9 injection) and 5 minutes-scan for ventral and dorsal views (for MyoRep mice). The analyses were made with the Living Image Software (Perkin Elmer).

### Mouse fasting and refeeding

Mice were subjected to fasting for either 16 or 48 hours. Fasted animals were transferred to new, clean cages without food but with free access to water, while maintaining environmental enrichment. After 48 hours of fasting, food was reintroduced. *In vivo* imaging was performed 16 or 48 hours after food deprivation and 24 hours after refeeding.

### Unilateral cut of the sciatic nerve of mice

Cutting the sciatic nerve was done on *wild-type* mice to induce atrophy of the hindlimb muscles. Anesthetized mice were placed in a prone position and a small incision was made on the thigh to expose the sciatic nerve. The nerve was isolated using tweezers, and a 2-3 mm segment was excised to prevent reinnervation. The incision was closed with absorbable sutures and disinfected with betadine. Sham-operated animals underwent the same procedure without nerve cutting.

### Mouse sacrifice and tissue collection

Mice have been euthanized by decapitation when sleeping because of exposure to gas anaesthesia with a mixture of 100% oxygen (2 l/min) and 5% isoflurane. Mice have been sacrificed when they reached a BWL higher than 20% for 3 consecutive days or when the tumor reached 10% of total BW (and before ulceration) or when four out of five of the following parameters were present in the same moment: immobility, hypothermia, kyphosis, tremor and ruffled coat. Apc^Min/+^ mice have been sacrificed through similar criteria, but also considering different time points (15 and 18 weeks of age). We collected blood to prepare plasma, hindlimb muscles (TA, GAS, soleus, quadriceps, back muscles). To prepare plasma, blood was collected from mice in BD Microtainer additive K2-EDTA beadless cap (BD, Milan, Italy), tubes that contain EDTA as anticoagulant, and centrifuged for 5 minutes at 5,000 x g at 4°C. Once dissected, muscles and organs have been weighed with analytical scale (Cabru S.a.s., Biassono, Italy) and frozen in liquid nitrogen-cooled isopentane (−190°C) or analysed for *ex vivo* imaging with IVIS system.

### Total RNA extraction from muscles

Total RNA was isolated from muscles with QIAzol Lysis Reagent and miRNeasy Kit (Qiagen, Hilden, Germany). The frozen muscle was cut perpendicularly to the tendon and resuspended in 700 μl of QIAzol (Qiagen, Hilden, Germany). Muscles were then homogenized with T25 digital Ultra-Turrax homogenizer (IKA). RNA was extracted following manufacturer’s instructions using chloroform and RNeasy Mini column provided by the kit. RNA concentration, purity and integrity were measured in a spectrophotometer (NANODROP 1000, ThermoFisher Scientific, Waltham, MA, USA).

### Reverse transcription PCR

For the reverse transcription of RNA into cDNA, High-Capacity cDNA Reverse transcription Kit (Applied Biosystems, Waltham, MA, USA) was used. One μg of RNA was added to 4 µl of buffer 10X, 1,6 µl of dNTPs, 4 µl of random primers, 2 µl of reverse transcriptase enzyme and water reaching a final volume of 40 µl. Then, samples were placed in the thermocycler (BioRad T100 Thermal Cycler) at 25°C for 10 minutes, 37°C for 2 hours and 85°C for 5 minutes.

### Quantitative Polymerase Chain Reaction

Analysis of mRNA in muscle was done using TaqMan reverse transcription reagents (Life Technologies Waltham, MA, USA) or the fluorescent intercalating DNA SYBR Green (Life Technologies Waltham, MA, USA). In each well of a Fast Optical 96 well-reaction plate, a mix containing TaqMan Master mix and the probe for TaqMan assay (Life Technologies Waltham, MA, USA) or SYBR Green Master mix and oligonucleotides as primers (Metabion, Germany) were added to 2 µl of cDNA (20 ng). Water was added to reach a final volume of 11 µl. The instrument we used for these assays was a 7900HT Fast Real-Time PCR System (ThermoFisher Scientific, Waltham, MA, USA). The program for Real-Time PCR was as follows: step 1, 95°C for 15 min; step 2, 95°C for 25 sec; step 3, 60°C for 1 min; repeating steps 2 and 3 for 40 cycles. *Gusb* (β-Glucuronidase) or *Tbp* (Tata binding protein) or *Ipo8* (Importin 8) were used as housekeeping genes. The following table contains the list of all primers used in this work.

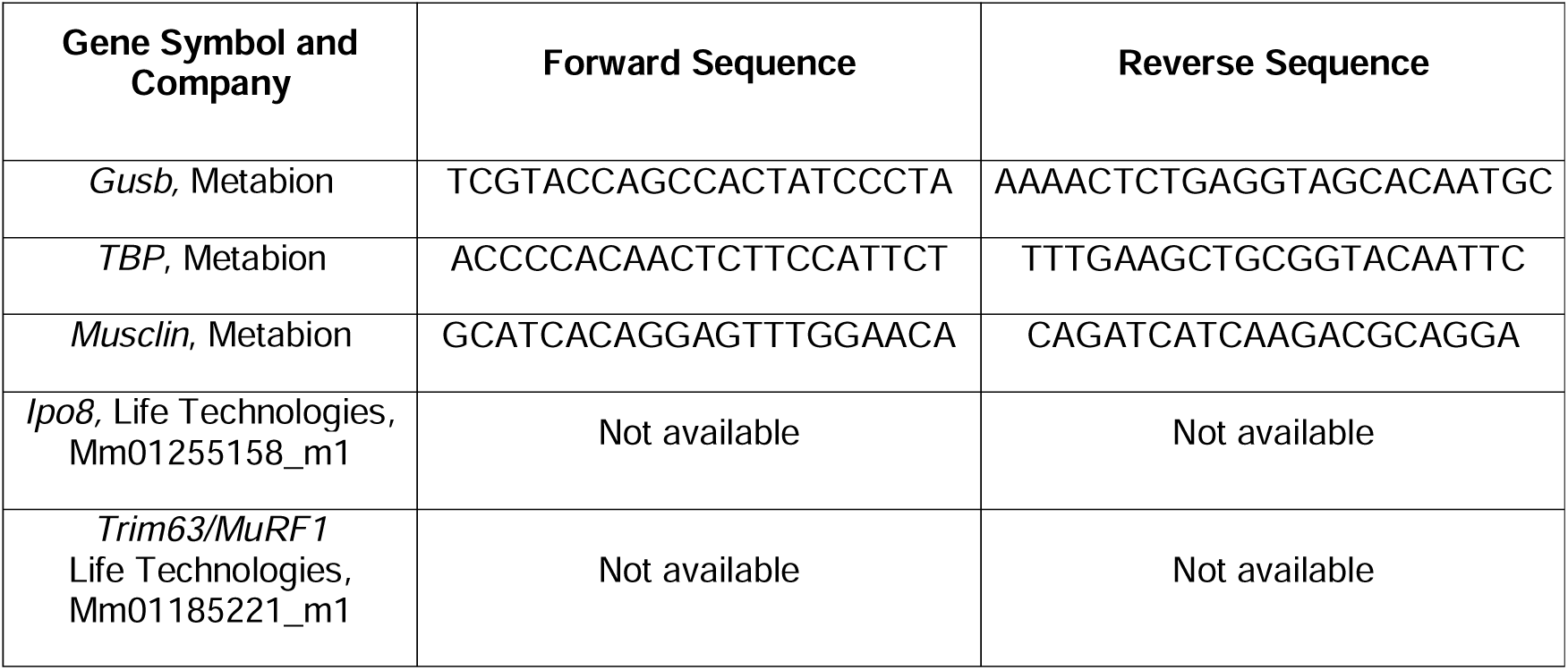

### Total protein extraction from tissues

Total proteins were extracted from muscles using radioimmunoprecipitation assay buffer (RIPA buffer) with the final addition of 4% sodium dodecyl sulphate (SDS), protease and phosphatase inhibitors (Roche, Basel, Switzerland). A volume of RIPA buffer corresponding to 20 times the weight of the muscle was added, then muscles were then homogenized with T25 digital Ultra-Turrax homogenizer (IKA). Each sample was vortexed for 1 minute, and then incubated for 40 minutes at 4°C on rotating wheel. For the last step, samples were centrifuged at 13,000 x g for 20 minutes at 4°C and the supernatant was collected and stored at −80°C.

### Protein quantitation

The final protein concentration was quantitated by the bicinchoninic acid or BCA (Pierce, Waltham, MA, USA) or with Bradford if proteins were extracted with PLB from Luciferase assays. For BCA, each sample was diluted in water 1:10, and 10 µl were placed in a well of a flat transparent 96-well-plate, in duplicate. Then, 200 µl of BCA kit were added. On the same plate, known concentration of bovine serum albumin (BSA) were added. After 30 min of incubation at 37°C, using a spectrophotometer at 562 nm of wavelength, the absorbances were measured. The quantitation of the proteins was than extrapolated from the BSA standard curve based on the absorbance detected. For Bradford assay, each sample was diluted in water 1:10, and 10 µl were placed in a well of a flat transparent 96-well-plate, in duplicate. Then, 200 µl of Bradford substrate were added. On the same plate, known concentration of bovine serum albumin (BSA) were added. The absorbances were measured using a spectrophotometer at 595 nm of wavelength. Then, 10 µg of proteins were added to Laemmli sample buffer 4X (Biorad, Hercules, CA, USA) supplemented with 10% β-mercaptoethanol (Sigma, St. Louis, MO, USA), and boiled at 97°C for 7 min.

### Western Blotting

Each sample is placed into a well of a precast and gradient (4-20%) gel of sodium dodecyl sulfate polyacrylamide (BioRad) for electrophoresis (SDS-PAGE). The run was performed soaking the gels in running buffer (10% TGE, 1% SDS and H_2_O) and applying at constant voltage (100 V) for 90 min. Later, proteins on gel were transferred to polyvinylidene difluoride membrane (PVDF; GE Healthcare, Chicago, IL, USA) using a transfer buffer (10% TGE, methanol and H_2_O), for 1 hour at constant amperage (400 mA). Then, the membrane was incubated at RT with a 5% BSA solution (BSA – bovine serum albumin - dissolved in Tris-Buffered Saline-Tween (TBS-T) 1%). Then, the membrane was incubated overnight with a solution with 5% of BSA containing the primary antibody directed to the protein of interest. We used the following primary antibodies: Anti-MuRF1, kindly donated by Prof. Alfred L. Goldberg (Harvard Medical School, Boston, MA, USA) and anti-vinculin (V9264, Sigma). After the overnight incubation with the primary antibody and a 30 min wash with TBS-T, the membrane was incubated for 1 hour RT with the secondary antibody, diluted in a solution of 1% BSA in TBS-T and conjugated with alkaline phosphatase. The membrane was then washed with TBS-T for 30 minutes and, using CDP-star substrate (ThermoFisher Scientific, Waltham, MA, USA), proteins were detected by means of a chemiluminescence reaction, capture by the Odyssey Imager (Li-Cor, Bad Homburg, Germany). The intensity of the band was analysed using the ImageJ software (National Institutes of Health, Bethesda, MA, USA).

### Enzyme-Linked Immunosorbent Assay

The levels of corticosteroids were measured in murine plasma of fed and fasted mice and PBS and 4T1-injected mice using an enzyme-linked immunosorbent assay (ELISA) kit (RTC002R, BioVendor, Brno, Czech Republic). Ten µl of plasma or standard curve were added to a well of the kit’s plate. After the addition of 100 µl of Incubation buffer and 50 µl of Enzyme conjugate, the plate was incubated for 2 hours at RT. After four washes, 200 μl of Substrate solution was added to each well and the plate incubated for 30 min in the dark. Lastly, after the addition of 50 µl of stop solution, the optical density absorbance was read at 450 nm. The detection range of the kit is 11.4-819 ng/ml, depending on the sex of the mouse and the time of blood sampling. The lowest analytical detectable level of corticosterone that can be distinguished from the Zero Calibrator is 6.1 ng/ml. Musclin levels in murine plasma were measured in an ELISA. Plasma was collected in 0.5 M EDTA, centrifuged 10 min at 10,000 rpm at 4°C, and then stored at −80°C. Musclin was quantitated with a Cusabio ELISA kit (CSB-EL017269MO, Tema Ricerca, Italy). Diluted plasma samples and standards were applied to the ELISA plates, and incubated for 2 hours at 37 °C. Plates were then incubated with biotinylated antibody for 1 hour at 37°C. After three washes, plates were incubated with horseradish peroxidase (HRP)-avidin 1 hour at 37°C, then washed and incubated with the tetramethylbenzidine substrate for 15 min at 37 °C. Optical density was detected at 540 nm. The detection range of the kit is 15.6–1000 pg/mL and the sensitivity is about 3.9 pg/mL.

### Statistical analysis

Sample size was determined by power analysis with G*Power on the basis of similar experiments previously published by our laboratory. All the experiments were repeated at least twice. For statistical analysis, data (means ± standard errors of the mean or SEMs) were analyzed with GraphPad Prism 10.2 for Windows (Graph-Pad Software, San Diego, CA, USA) with the following statistical tests: ordinary one-way analysis of variance (ANOVA) for multiple comparisons followed by Tukey’s or Dunnett’s post hoc test, Kruskal-Wallis test followed by Dunn’s post hoc test, two-way ANOVA for multiple comparisons followed by Tukey’s post hoc test, unpaired t-test or Mann-Whitney test for comparisons of two groups, paired t-test test for paired analysis, Brown-Forsythe test for variance; * p≤0.05; ** p≤0.01; *** p≤0.001; **** p≤0.0001.

## RESULTS

### *In vitro* tests to identify the best early reporter sequence to sense myoblast atrophy

Throughout the years, we have been testing a long list of sequences both encoding for binding sites for transcription factors known to play a role in atrophy as FoXO3, NF-kβ ^7^, STAT3 ^8^, Smad 2/3 ^9^ or promoters of well-known atrogenes (atrogin-1 and MuRF1). Most of these sequences were provided in Firefly Luciferase reporter plasmids by collaborators (see Materials and Methods) or engineered by us and listed in (**Table I**).

These vectors were transfected for 24 hours *in vitro* in C2C12 myoblasts and their possible response to various atrophic stimuli (supernatants of cancerous cells or HBSS to mimic starvation or dexamethasone) was measured to see if anticipated protein loss. These experiments enabled us to understand that MuRF1 promoter was the best. The original length or MuRF1 promoter is 5 Kb, too long to be used to generate a reporter *in vivo*. So, we generated the so-called murine MuRF1 that is 628 bp long and was selected as minimal one to signal atrophy *in vitro*. Nonetheless, we generated two more variants from such mMuRF1 to test if Twist transcription factor binding sites and glucocorticoid-responsive elements or GRE were needed or not to improve the earlier response to atrophy and its specificity. So, we compared *in vitro* the murine promoter of MuRF1 (or simply mMuRF1 that does not display changes to the genomic sequences), with mMuRF1 TWIST (or simply TWIST), and mMuRF1 TWIST GREDEL (or simply GREDEL). All these promoters were cloned upstream of Firefly Luciferase and co-transfection with a TK-Renilla Luciferase vector served as control to normalize the data.

As example, we transfected C2C12 myoblasts with each of these 3 vectors for 24 hours and exposed them to media conditioned from C26 cells, as atrophying stimulus, or from 4T1 cells as non-cachectic one or from C2C12 cells and DMEM as controls for the next 24 hours. We found that C26 supernatant induces the activity of Firefly Luciferase, measured in Luciferase-based assay and normalized over Renilla Luciferase for all three vectors compared to C2C12 supernatant and DMEM. Importantly, only TWIST and GREDEL were able to discriminate between the non-cachectic 4T1 and the cachectic C26, indicating that mMuRF1 promoter was less specific to the atrophic stimulus (**Figures 1A-C**).

**Figure 1.**
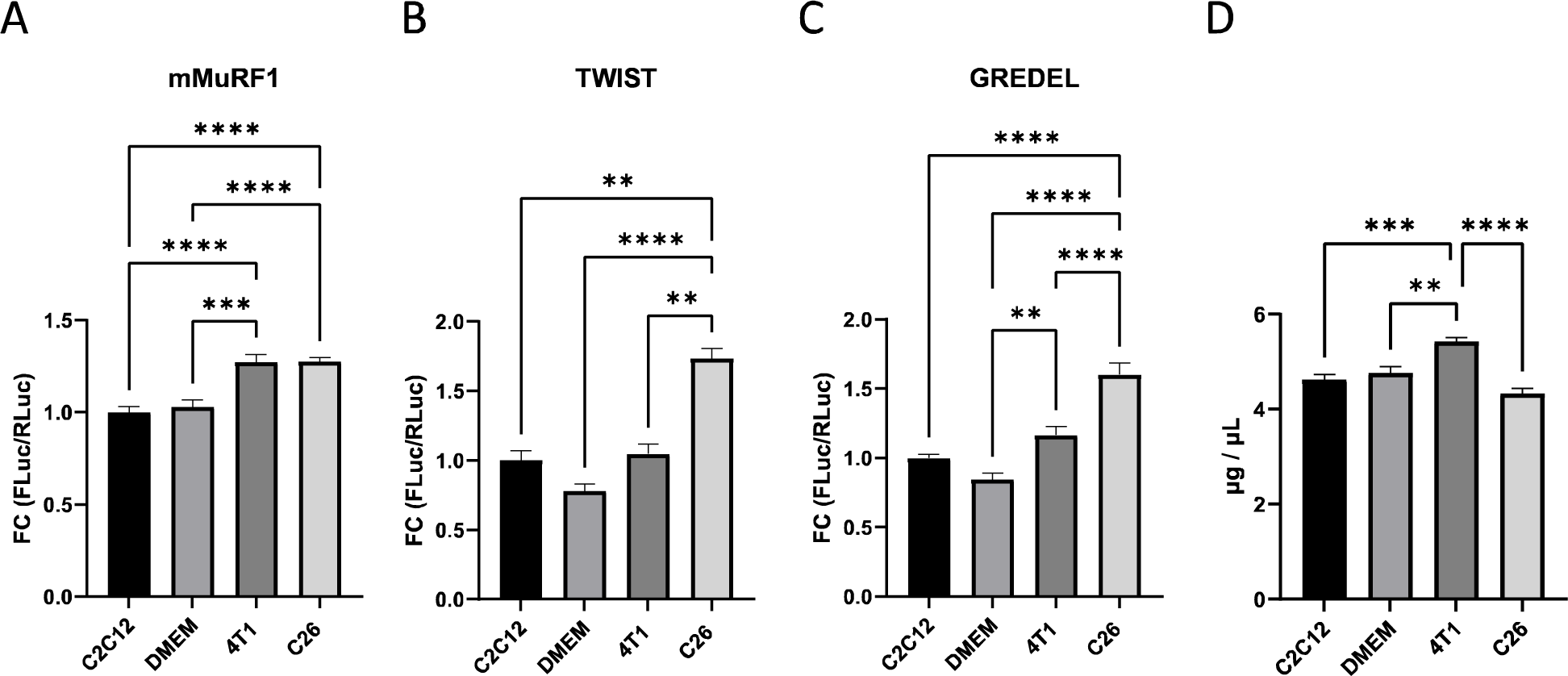
The variants TWIST and GREDEL of the murine promoter of MuRF1 (mMuRF1) can discriminate between media conditioned by cachectic from non-cachectic cell lines, even before protein loss. Luciferase assays of C2C12 myoblasts transfected with mMuRF1 (A), TWIST (B) and GREDEL (C) reporter plasmids (all co-trasfected with pRL-TK-Renilla to normalize the data) for 24 hours and treated with conditioned media from C2C12, 4T1 and C26 cell lines or DMEM as control for the next 24 hours. One-way ANOVA or Kruskal-Wallis’s test followed by Dunnett’s or Dunn’s post hoc test. **p≤0.01, ***p≤0.001, ****p≤0.0001. N = 12. (D) Protein content of samples from (A-C) was analyzed by Bradford assay. One-way ANOVA followed by Tukey’s post hoc test. **p≤0.01, ***p≤0.001, ****p≤0.0001. N = 6. All data are reported as mean ± SEM.

To further test whether these vectors were able to signal atrophy before proteins diminished, we measured the protein content on the same samples whose luciferase activities are shown in **Figures 1A-C**. Aside from a slight protein accumulation in cells exposed to 4T1 conditioned medium (**Figure 1D**), our data indicate that overall, the variants TWIST and GREDEL of the murine promoter of MuRF1 can drive *Firefly Luciferase* expression upon cachectic stimulus, anticipating protein loss.

### GREDEL reporter is better than TWIST to sense *in vivo* systemic or local pathological atrophy

We next compared the TWIST and GREDEL vectors by electroporating them in TA of MCG101-bearing mice as model of cachexia for their ability to signal *in vivo* atrophy. This model shows body weight loss (BWL) 18 days after tumor injection compared to PBS-injected mice (**Figure 2A**). When we sacrificed mice at 6-7, 14-15 and 20 days after tumor cell injection, we found that tumor weight was significantly greater at 14-15 and 20 days compared to 7 days (**Figure 2B**). TA and GAS weight were reduced in size only at 20 days following tumor injection compared to 6-7 days and to PBS-injected mice (**Figure 2C and Supplementary Figure 1A**). Instead, MuRF1 mRNA expression was already increased in TA at 14-15 days, anticipating muscle wasting (**Figure 2D**).

**Figure 2.**
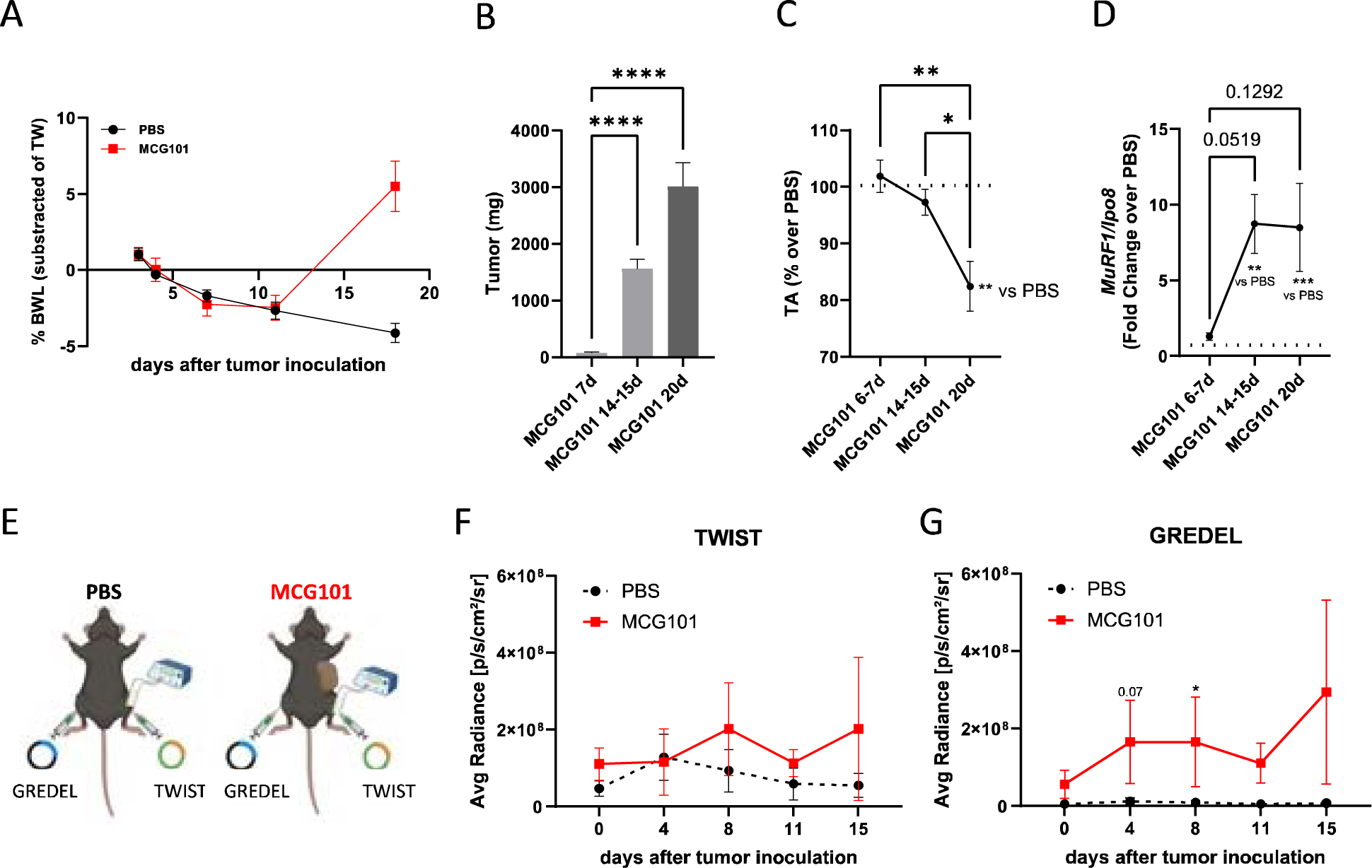
MCG101-carrying mice display atrophy detectable earlier by GREDEL than TWIST reporter. (**A**) Percentage of body weight loss (BWL%) subtracted of tumor weight of C57BL6 male mice injected subcutaneously with PBS or MCG101 is plotted over time. Two-way ANOVA followed by Tukey’s post hoc test. ****p≤0.0001. N = 5. (**B**) Tumor weight of MCG101 hosts sacrificed at different times. Kruskal-Wallis test followed by Dunn’s post hoc test. ****p≤0.0001. N = 7-33. (**C**) Tibialis anterior (TA) weight (% over PBS, dotted line) of MCG101 carriers sacrificed at different times. One-way ANOVA followed by Tukey’s post hoc test. *p≤0.05, **p≤0.01. MGC101 20d vs PBS, Unpaired t test, **p≤0.01. N = 7-33. (**D**) MuRF1 mRNA expression measured by qPCR in TA from mice of panel **C**. One-way ANOVA followed by Tukey’s post hoc test and unpaired t test. **p≤0.01, ***p≤0.001. (**E**) Graphical representation of the experimental scheme, made with Biorender. Analysis of photon emission of TA expressing TWIST (**F**) or GREDEL reporters (**G**) comparing the area under the curve (AUC) of PBS and MCG101 hosts. (**F**) Unpaired t test for AUC, *p≤0.05, N=3-5; (**G**) Unpaired t test for AUC, **p≤0.01, N=3-5. At days 4 and 8 PBS vs MCG101: *p≤0.05, Mann-Whitney test. All data are reported as mean ± SEM.

TWIST- and GREDEL-expressing plasmids were electroporated in TA from MCG101- and PBS-bearing mice, as shown in **Figure 2E**. When we analysed the photon emission of TA expressing TWIST or GREDEL reporters, comparing the area under the curve (AUC) of PBS and MCG101 hosts, we found a greater bioluminescence emission in MCG101-bearing mice for both vectors within 15 days from tumor injection, before muscle atrophy (**Figures 2F and G**). Activation of both vectors was confirmed in TA from MCG101 carriers in *ex vivo* luciferase assay (**Supplementary Figure 1B**). Notably, GREDEL, but not TWIST, was activated already at 4 and 8 days after tumor injection in MCG101 hosts with respect to PBS ones (**Figure 2G**). TWIST showed greater variations in photon emissions with respect to GREDEL, when comparing the AUC of tumor-free mice, possibly indicating that TWIST is more sensitive than GREDEL to circadian cycle-related variations (**Supplementary Figure 1C**).

To validate the ability of TWIST and GREDEL reporters to detect local muscle atrophy, we further tested them in a mouse model where cut of the sciatic nerve induces atrophy of GAS and TA muscles ^18^. The denervated leg showed loss of the TA and GAS by 15-20% only 7 days after surgery and by 30-50% 14 days after denervation, compared to the sham-operated leg (**Figure 3A**). Interestingly, protein levels of MuRF1 were increased already 3 days after surgery in denervated TA (**Figures 3B and C**), anticipating muscle atrophy, as shown by others in ^6^. To assess the bioluminescent signal, each mouse was electroporated two weeks before denervation in both TAs with either TWIST- or GREDEL-encoding plasmids, and the denervated leg was compared to the sham-operated one. Interestingly, while TWIST reporter got activated 7 days after denervation (**Figure 3D**), GREDEL was able to drive Firefly Luciferase activity as early as 1–2 days in the denervated leg compared to the sham-operated one (**Figure 3E**), anticipating muscle mass loss.

**Figure 3.**
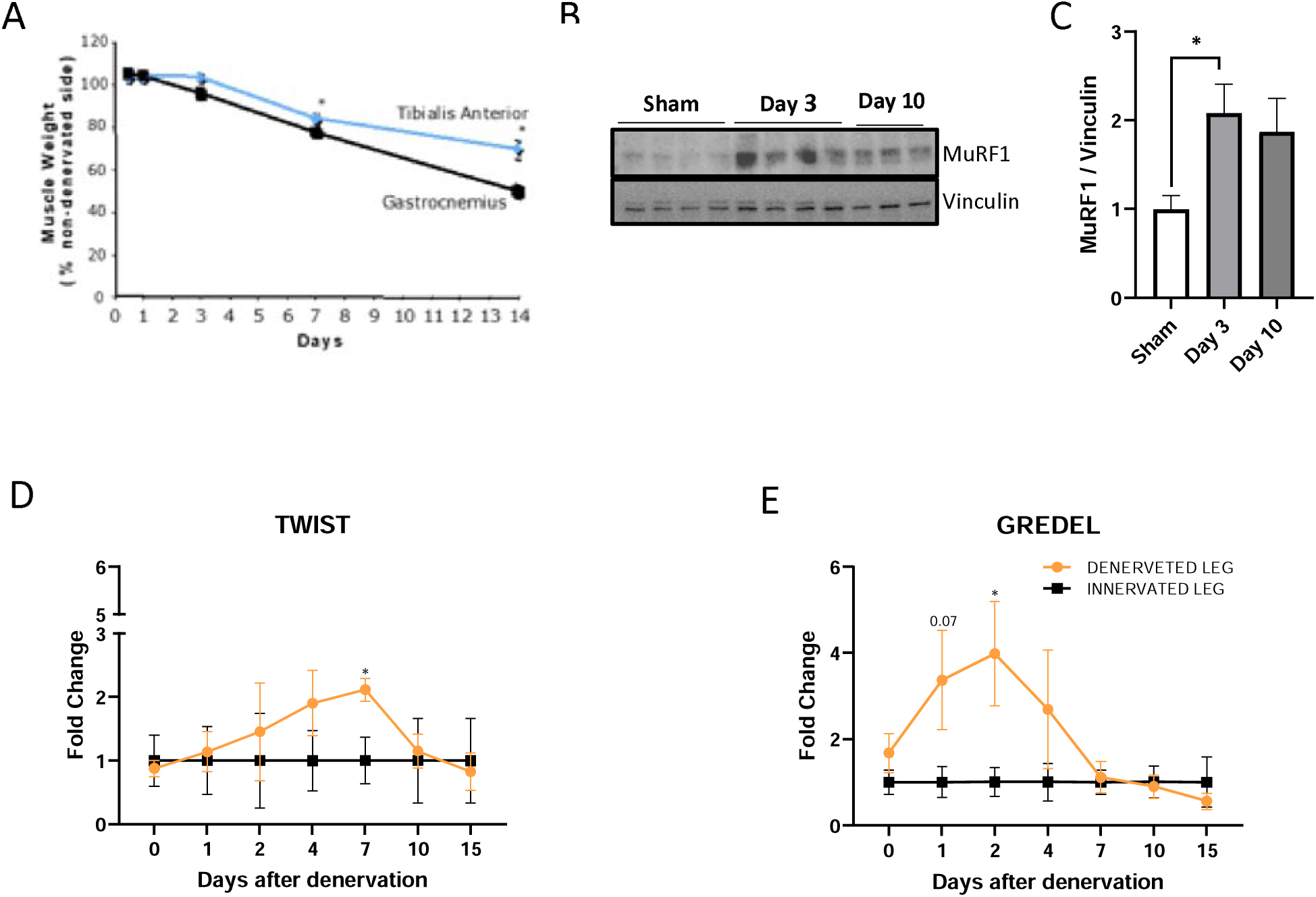
GREDEL reporter detects earlier atrophy also in mice subjected to denervation. (**A**) Gastrocnemius (GAS) and tibialis anterior (TA) weight (% of non-denervated side) of denervated leg at different times. One-way ANOVA test followed by Dunnett’s post hoc test. *p≤0.05. N = 3. (**B**) Western blot for MuRF1 of TA of sham-operated leg and denervated leg at day 3 and 10 after cutting the sciatic nerve. Vinculin is used as loading control. (**C**) Quantitation of western blot in **B**. Unpaired t test. *p≤0.05. N = 3-4. Analysis of photon emission of TA expressing TWIST (**D**) or GREDEL reporters (**E**) upon denervation. Fold change of denervated leg over innervated one is shown. (**D**) Two-way ANOVA followed by Tukey’s post hoc test. *p≤0.05, N=4. (**E**) Two-way ANOVA followed by Tukey’s post hoc test. Unpaired t test for days 1 and 2. *p≤0.05, N=4. All data are reported as mean ± SEM.

### TWIST reporter is more undesirably sensitive than GREDEL to *in vivo* physiological atrophy

Glucocorticoids are known to induce muscle atrophy both *in vitro* and *in vivo* ^19^. They contribute to muscle atrophy in cancer cachexia ^20^ and denervation ^21^, but they are also released from the hypothalamic-pituitary-adrenal axis during fasting in mice ^22^. Moreover, corticosterone and cortisol are secreted in a circadian-dependent manner in both animals and humans ^23,24^, being at the intersection between pathological and physiological atrophies.

Anorexia occurs also during cancer cachexia, but we aimed to generate a reporter mouse sensitive only to cachexia and not to the possible reduced food intake. Even if MCG101 hosts eat as well as PBS mice (**Supplementary Figure 2**), other cachectic tumors cause also anorexia in mice, as C26 ^25^. These considerations prompted us to compare TWIST and GREDEL reporters also in response to physiological atrophies induced by circadian rhythms or fasting. The best reporter to pathological atrophies should be insensitive to either of them.

Firstly, we confirmed in Luciferase assays that only C2C12 transfected with mMuRF1 and to a lesser extent with TWIST, but not with GREDEL reporter plasmids responded to 1 or 10 µM dexamethasone (Dexa) (**Figures 4A-C**). This occurred when protein level was not yet affected by Dexa treatment (**Figure 4D**). Secondly, when we electroporated these vectors in TA and subjected mice to 48 hours of fasting, unlike GREDEL, TWIST was activated (**Figure 4E**) and its emission reduced upon refeeding, confirming its undesirably sensitivity to a physiological type of atrophy. Additionally, despite not significant, GREDEL exhibited a trend toward lower variability than TWIST in the bioluminescence signal detected in the TA of healthy mice between morning and afternoon (**Figure 4F**), in accordance with data showed in **Supplementary Figure 1C**.

**Figure 4.**
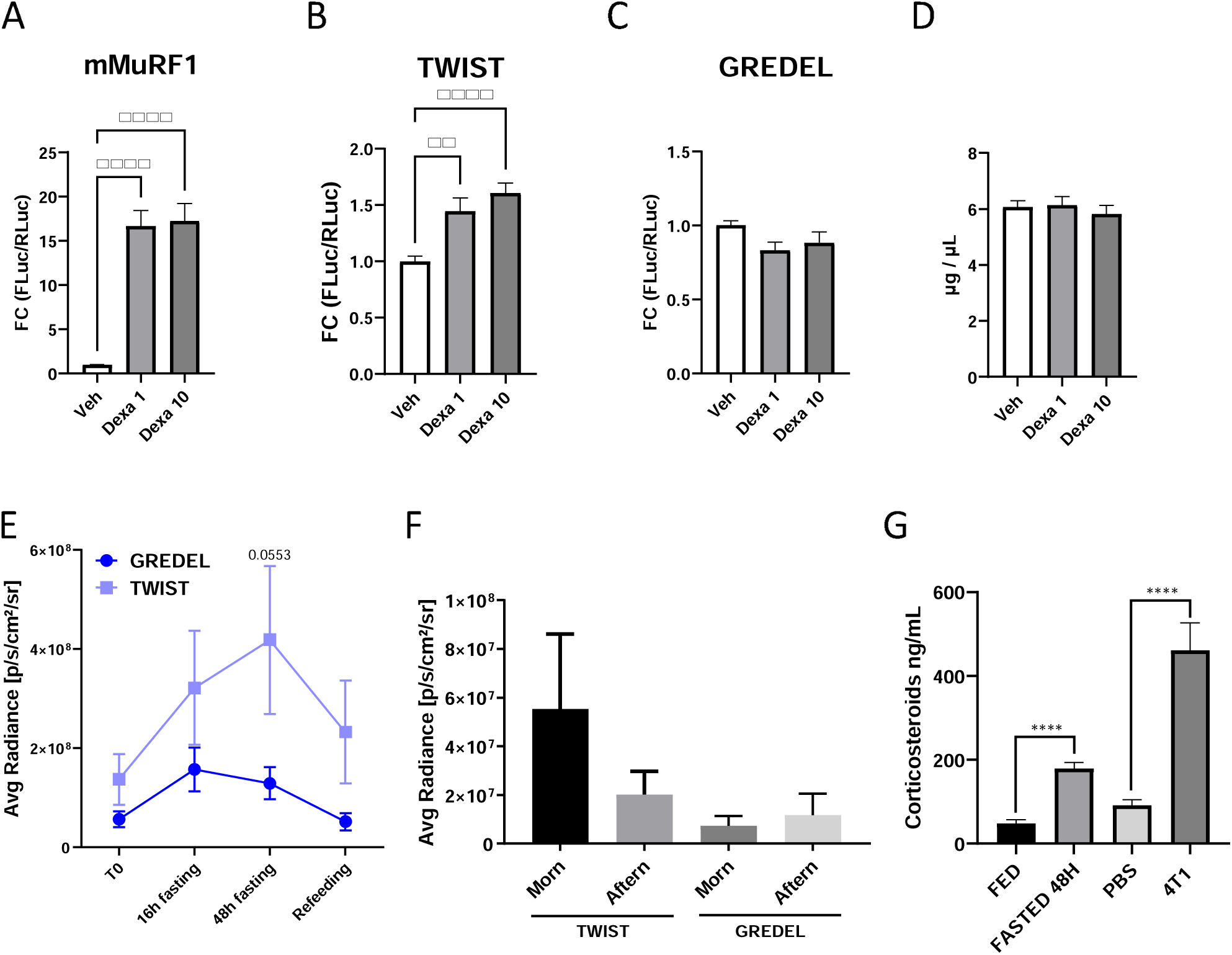
GREDEL reporter is not sensitive to circadian rhythms variations as TWIST. Luciferase assay analysis of C2C12 myoblasts transfected with mMuRF1 (**A**), TWIST (**B**) or GREDEL (**C**) reporter plasmids and treated with 1 or 10 µM dexamethasone (Dexa 1 oe 10, respectively). Kruskal-Wallis’s test followed by Dunn’s post hoc test. **p≤0.01, ****p≤0.0001. N = 14-23. (**D**) Protein content of experiments shown in (**A-C**) analyzed by Bradford assay. Kruskal-Wallis’s test followed by Dunn’s post hoc test. N = 12. (**E**) Analysis of photon emission comparing area under the curve (AUC) of TA expressing GREDEL reporter with TA expressing Twist reporter. Unpaired t test for AUC, *p≤0.05, N=12. (**F**) Analysis of photon emission comparing TA expressing GREDEL reporter with TA expressing Twist reporter in the morning and in the afternoon or tumor-free mice. One-way ANOVA and Brown-Forsythe test, not-significant. N=5. (**G**) ELISA for corticosteroids in plasma of mice fed or fasted or injected with PBS or 4T1 cells for 21 days. Unpaired t test, ****p≤0.0001, N = 5-10. All data are reported as mean ± SEM.

The activation of TWIST in response to fasting may be due to corticosteroids, as we observed higher plasma levels of corticosteroids in mice fasted for 48 hours compared to fed mice (**Figure 4G**). Of note, corticosteroid levels were also found elevated in plasma from 4T1-bearing mice compared to PBS-injected controls (**Figure 4G**). 4T1 is a triple negative breast cancer cell line that does not cause cachexia *in vivo* ^26^. Altogether, these data prompted us to choose to generate AAV9 and then the reporter mouse with the GREDEL promoter and to rename it MyoRep.

### MyoRep discriminates sex-specific muscle atrophy in cachectic Apc^Min/+^ mice

Then, we generated an AAV9 expressing MyoRep upstream of *Firefly Luciferase*. To further validate its functionality during cancer cachexia, we injected MyoRep-AAV9 into the TA muscle of Apc^Min/+^ and WT mice of both sexes at 8 weeks of age. We then monitored the mice using *in vivo* imaging until 18–19 weeks of age, when muscle atrophy became evident ^27^. The Apc^Min/+^ mouse model develops spontaneously intestinal tumors and cachexia, mimicking colorectal cancer-associated muscle wasting with males more severely affected than females ^28, 29, 30, 31^.

Once a week, we imaged the mice under anaesthesia using IVIS machine (Pekin Elmer) and tracked the onset and progression of muscle atrophy by measuring the bioluminescent signal from the electroporated leg (**Figure 5A**). The region of interest (ROI) was quantified using dedicated software, and the signal was plotted over time (**Figure 5B**). As expected, Apc^Min/+^ males exhibited a higher signal than age-matched Apc^Min/+^ females, starting from 15 weeks of age (**Figure 5B**). Additionally, we weighed the mice weekly and plotted their body weights over time, further confirming the sex differences in body weight loss (BWL), starting from 16 weeks of age (**Figure 5C**). This indicates that MyoRep activation precedes BWL by one week in these cachectic mice. Interestingly, sex-specific activation of MyoRep is also detected in TA collected at 15 weeks of age and analysed in *ex vivo* Luciferase Assay (**Figure 5D**), confirming the data obtained *in vivo*. Moreover, MyoRep analysis proved to be more sensitive and less variable in detecting muscle atrophy *in vivo* compared to measuring MuRF1 *ex vivo* by western blot in TA muscle isolated from mice at 15 weeks of age (**Figures 5E and F**). In fact, MyoRep vector does not contain just MuRF1 promoter, but also repeated and appropriately modified sequences of MuRF1 promoter designed to enhance sensitivity to muscle atrophy while also removing GREs, as previously described.

**Figure 5.**
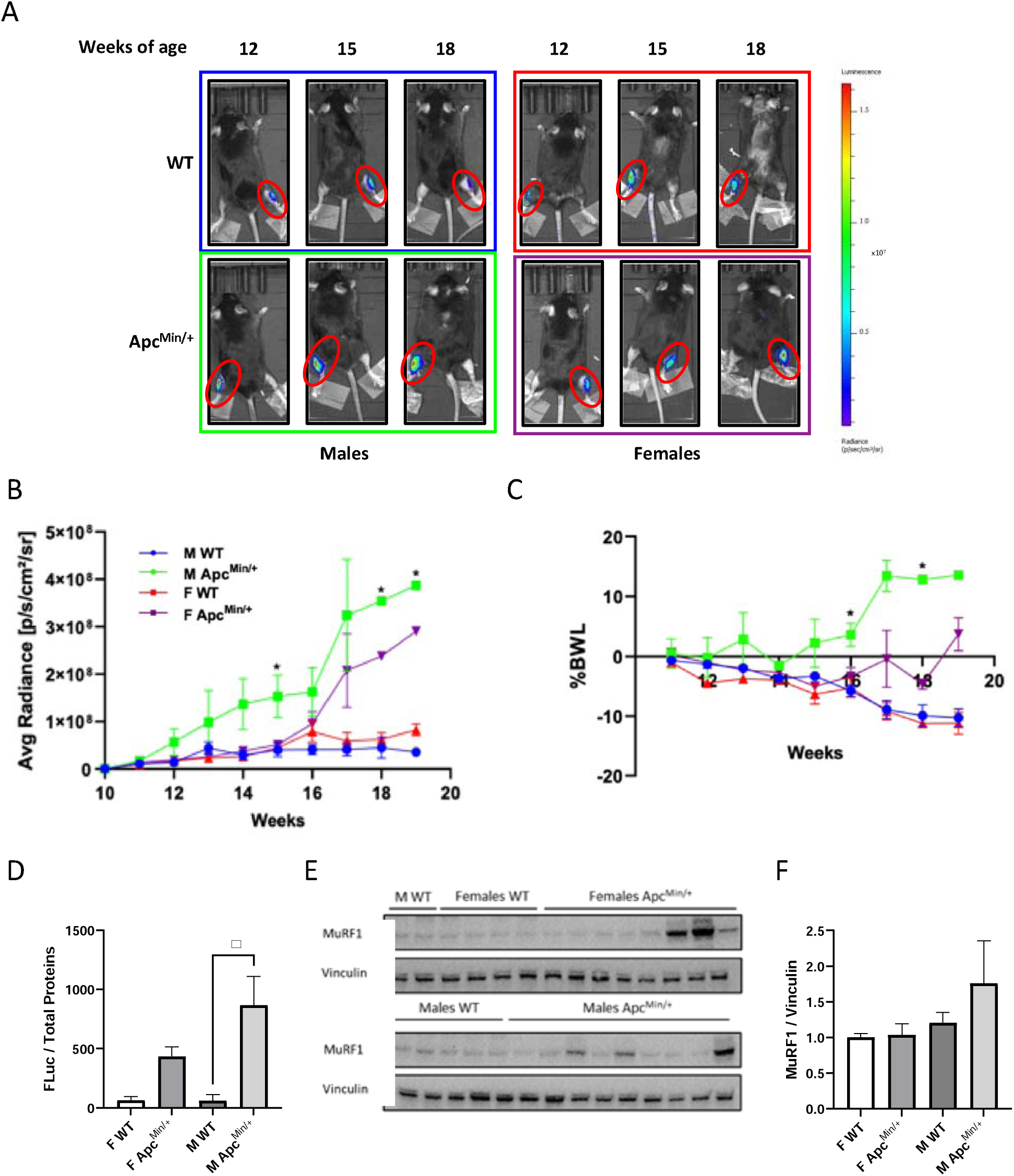
The GREDEL reporter can detect the sex-specific atrophy of cachectic Apc^Min/+^ mice. (**A**) Representative images of C57BL6 wt and Apc^Min/+^ mice of both sexes acquired by in vivo imaging. One TA was injected at 8 weeks of age with 10^12 vg/mL of MyoRep-carrying AAV9, the contralateral leg was injected with PBS. (**B**) Analysis of photon emission from the region of interest over time (ROI on TA). Two-way ANOVA, Tukey’s post hoc test, *p≤0.05, N=4. (**C**) Percentage of BWL of C57BL6 WT and Apc^Min/+^ mice is plotted over time. Two-way ANOVA followed by Tukey’s post hoc test. *p≤0.05. N = 5. (**D**) Luciferase assay analysis of TA from male and female WT and Apc^Min/+^ mice at 15 weeks of age. Total protein content was used to normalize. One-way ANOVA, Tukey’s post hoc test, *p≤0.05, N=3. (**E**) Western blot for MuRF1 of TA from male and female WT and Apc^Min/+^ mice at 15 weeks of age. Vinculin was used as loading control. (**F**) Quantitation of western blot in **E**. N = 4-9. All data are reported as mean ± SEM.

These data prompted us to use the GREDEL vector to generate the MyoRep reporter mouse, which expresses this vector throughout the skeletal muscles and heart (**Supplementary Figure 3**). This makes it a valuable tool for studying virtually any disease with associated muscle atrophy and *MuRF1* induction.

### The MyoRep mouse detects early systemic and local atrophy in *in vivo* imaging

To assess MyoRep’s ability to detect muscle atrophy, we used the MCG101 mouse model of cancer cachexia, as previously shown in **Figure 2**. Following the subcutaneous injection of MCG101 cells, we monitored mice using *in vivo* imaging over time until day 20, imaging them 6, 17 and 20 days and analysing photon emission from the region of interest (ROI) in the leg, dorsal and ventral views (**Figures 6A-C**).

**Figure 6.**
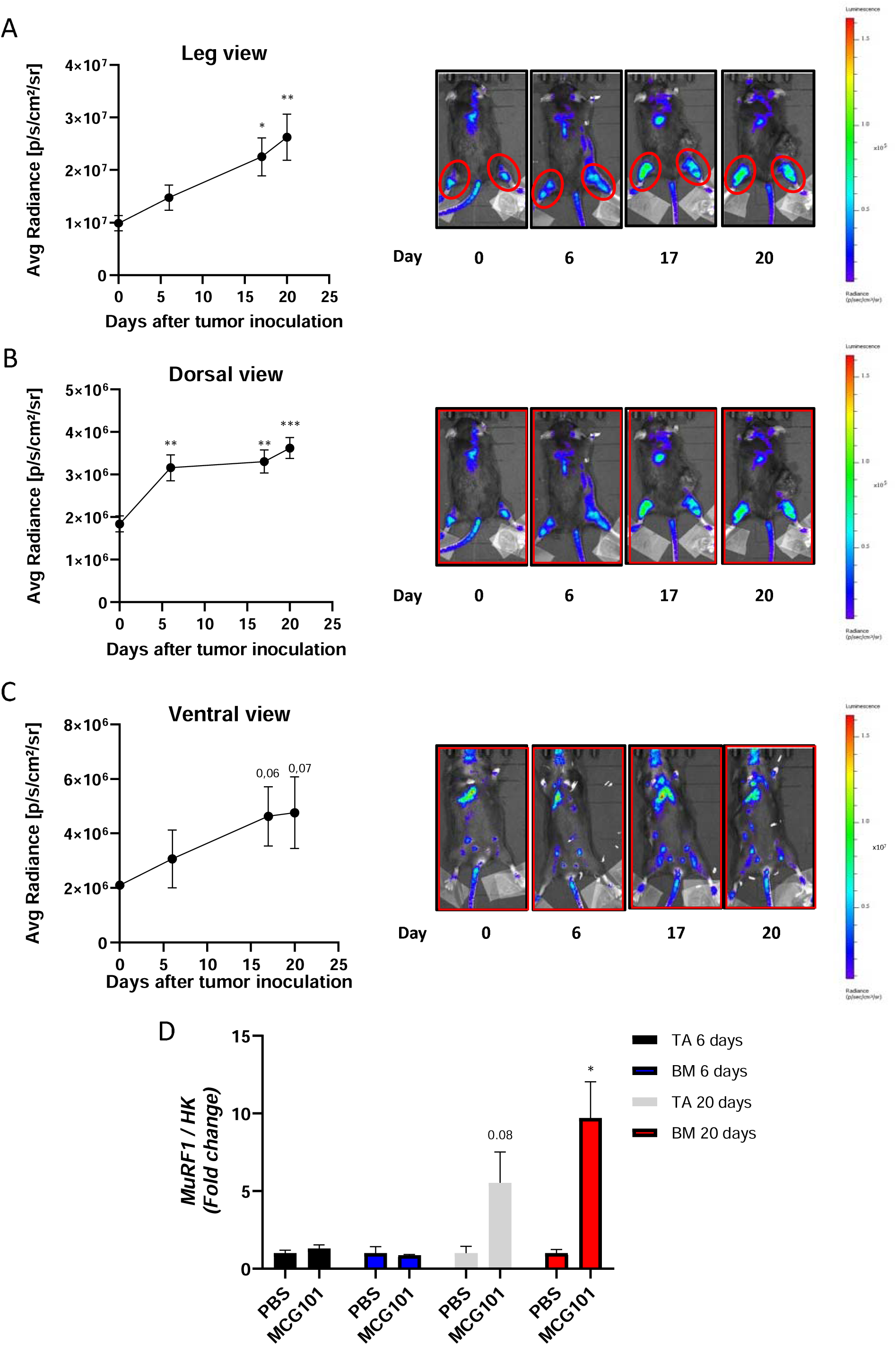
The dorsal scan of MCG101-carrying MyoRep mice detects earlier atrophy than leg or ventral views. (**A**) Analysis of photon emission from the region of interest over time (ROI, leg in red) and representative images of MCG101 bearing-MyoRep mice acquired by *in vivo* imaging. One-way ANOVA, Dunnett’s post hoc test vs day 0, *p≤0.05, **p≤0.01, N = 5. (**B,C**) Analysis of photon emission from the ROI in red on whole mouse on dorsal view (**B**) or ventral view (**C**) and representative images of MCG101 bearing-MyoRep mice acquired by *in vivo* imaging. (**B**) One-way ANOVA, Dunnett’s post hoc test vs day 0, **p≤0.01, ***p≤0.001, N = 5. (**C**) One-way ANOVA, Dunnett’s post hoc test vs day 0, N = 5. (**D**) *MuRF1* expression measured by qPCR in TA and back muscle (BM) 6 and 20 days after tumor inoculation. HK = housekeeping gene for normalization (β-Glucuronidase, *Gusb*, or Tata binding protein, *Tbp* or Importin 8, *Ipo8*). Unpaired t test. *p≤0.05, N = 3-4. All data are reported as mean ± SEM.

The bioluminescence emission analysis showed that MyoRep was activated at days 17 and 20 after tumor injection in the legs and ventral total view (**Figures 6A and C**), and as early as day 6 in the dorsal total view (**Figure 6B**) – at times when muscle atrophy is not present yet (**Figure 2C**). This demonstrates the early activation of MyoRep that even anticipates increases of mRNA levels of *MuRF1*, as measured by qPCR in TA and back muscle, where *MuRF1* increases only at 20 days in the dissected tissues (**Figure 6D**).

To test if MyoRep transgenic mice detect also local atrophy, we denervated them as described in **Figure 3** and performed *in vivo* imaging analysis from day 1 post-denervation until day 35, as shown in **Figure 7A**. MyoRep mice subjected to denervation exhibited increased bioluminescence in the denervated leg compared to the sham-operated one, even at day 1 post-denervation (**Figure 7B**), predicting the onset of muscle weight loss (**Figure 3A**). At 35 days after denervation, we sacrificed mice collecting various tissues (muscles, spleen, brain, heart and liver) and we *ex vivo* imaged muscles and organs (**Supplementary Figure 4A**). Analysis of photon emission from dissected muscles from MyoRep mice, comparing the denervated leg with the sham-operated contralateral one, revealed bioluminescence activation in the TA, GAS, and soleus (**Supplementary Figures 4B-D**). As expected, no activation was observed in the quadriceps, as it was not affected by cut of the sciatic nerve, nor in the other organs analysed.

**Figure 7.**
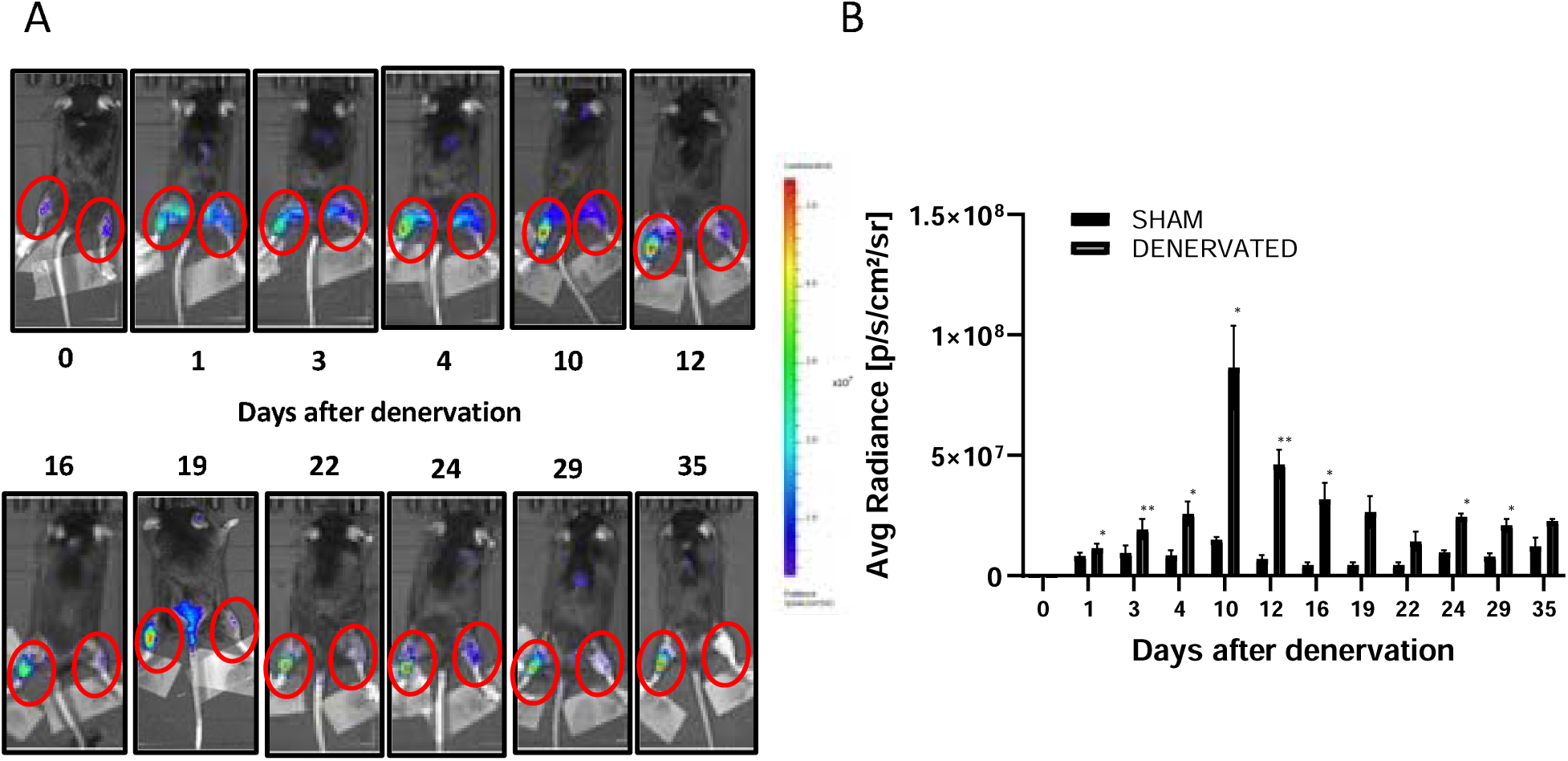
MyoRep mice subjected to denervation show an increased bioluminescent signal from the denervated leg before muscle weight loss. (**A**) Representative images of MyoRep mice acquired by *in vivo* imaging at different times. The left leg is denervated, while the right one is sham-operated. (**B**) Analysis of photon emission from the region of interest (ROI, hindlimb muscles, red circle). Paired t test, *p≤0.05; **p≤0.01. N = 4. All data are reported as mean ± SEM.

### The MyoRep mouse is insensitive to circadian rhythms and useful to *in vivo* evaluate anti-atrophic effects of molecules as musclin

As further control, we aimed to assess whether MyoRep mice were unresponsive to physiological muscle atrophy, as shown in **Figure 4**. We performed *in vivo* imaging in healthy mice in the morning and afternoon, as well as after 16, 24 or 48 hours of fasting, and observed no significant variations in photon emission across different ROIs (leg, dorsal, and ventral views) (**Figures 8A-C**). Importantly, these results confirm that our reporter mouse is not affected by circadian rhythms or fasting, maintaining its specificity for pathological muscle atrophy only.

**Figure 8.**
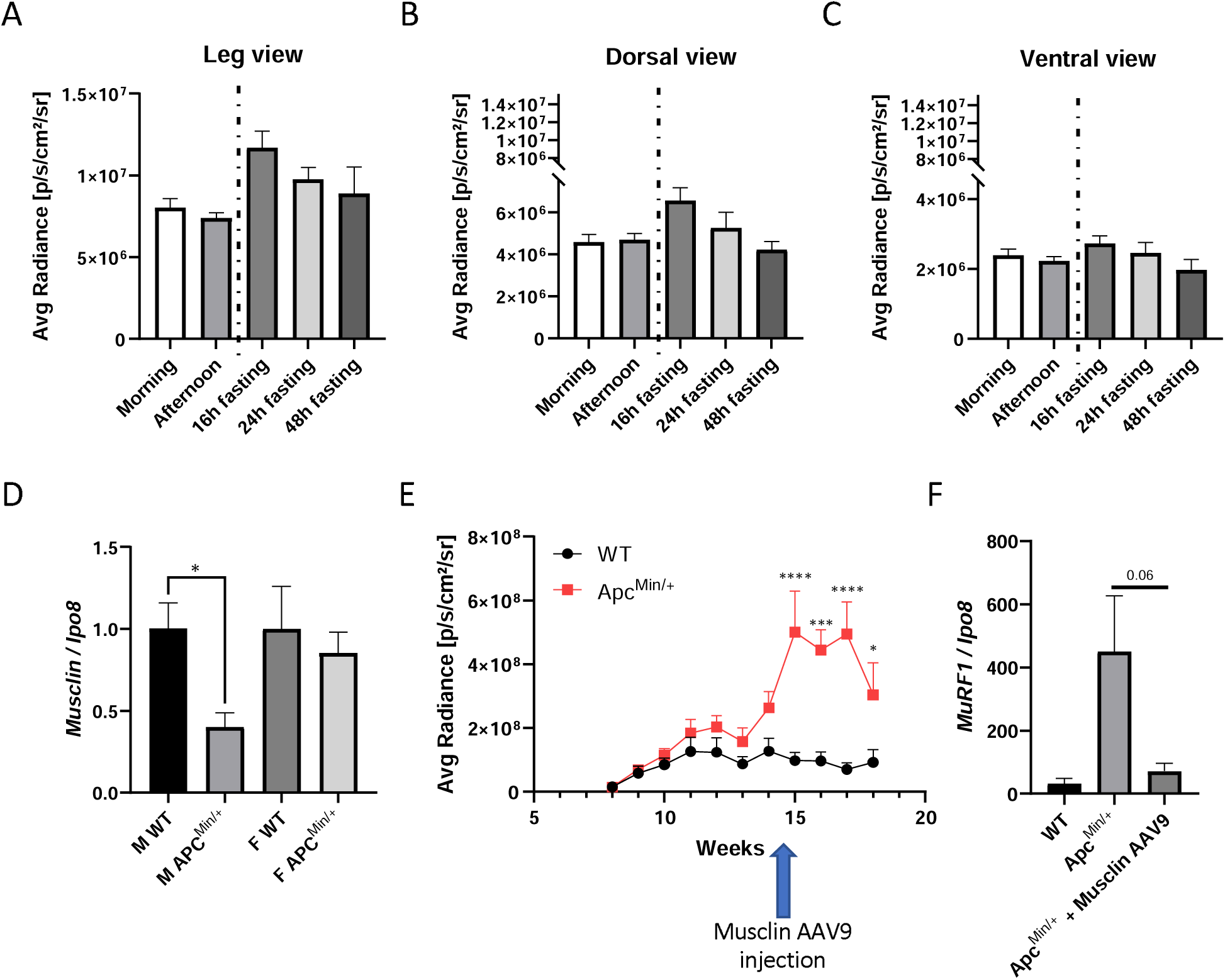
MyoRep mice do not display photon emission based on physiological muscle atrophy and decrease tumor-induced signal upon musclin-carrying AAV9 injections. (**A-C**) Analysis of photon emission from the region of interest (legs in **A**, whole mice in dorsal view in **B** and in ventral view in **C**) collected during different times of the day or with fasting of various lengths of MyoRep mice. One-way ANOVA, Tukey’s post hoc test. N = 4-5. (**D**) The mRNA expression of *musclin* measured by qPCR in TA of WT and APC^Min/+^ male and female mice at 12 weeks of age. *Ipo8* was used as housekeeping gene. Unpaired t test or Mann-whitney test. N = 7-13. *p≤0.01. (**E**) Analysis of photon emission from the region of interest over time (ROI on hindlimb leg). The arrow indicates the time of treatment in TA with musclin-AAV9. Two-way ANOVA, Tukey’s post hoc test, *p≤0.05, ***p≤0.001, ****p≤0.0001 N=5-7. (**F**) The mRNA expression of *MuRF1* measured by qPCR in TA after musclin-AAV9 injections. *Ipo8* was used as housekeeping gene. Unpaired t test. N = 3-5. All data are reported as mean ± SEM.

Finally, in the attempt to test whether MyoRep activation could be reversed by a possible anti-cachectic treatment, we measured musclin expression in plasma and muscles of Apc^Min/+^ mice at various ages. This is a myokine, in other words, a molecule released by muscles in response to exercise, whose levels we previously found to be reduced in TA of C26-bearing male mice and the electroporation of a plasmid encoding for musclin in TA of them partially preserved their fiber area ^32^. Interestingly, we found decreased levels of musclin mRNA in TA only in Apc^Min/+^ males and not in Apc^Min/+^ females at 12 weeks of age (**Figure 8D**), and also in the plasma at 18 weeks of age only in males (**Supplementary Figure 5**). So, we injected AAV9 expressing musclin into the TA of 14.5 week-old Apc^Min/+^ males and found that this treatment reduced photon emission at 18 weeks of age, restraining MyoRep activation (**Figure 8E**). In the same animals, we also measured MuRF1 levels in the TA at the time of sacrifice (18 weeks of age) to see whether musclin overexpression protected muscles from atrophy. We observed a trend toward reduced *MuRF1* mRNA levels, confirming that musclin acts as an anti-catabolic myokine and supporting MyoRep as able to detect these changes *in vivo* before sacrifice (**Figure 8F**). Of note, the binding sites inserted in the MyoRep are well conserved across species from *Ovis Aries* and *Sus Scrofa Domesticus* to *Homo Sapiens* (analysis done using BLAST database, **Supplementary Figure 6**), thus facilitating the application of MyoRep technology to other species.

These results highlight the potential of MyoRep to test drugs as able to mitigate muscle atrophy, as musclin.

## DISCUSSION

### Advantages of MyoRep technology over existing reporter tools

Muscle wasting is an unsolved medical issue that afflicts patients with very diverse chronic illnesses, causing premature death. To find novel *in vivo* tools to study muscle atrophy should be prioritized to accelerate the discovery of biomarkers predicting in advance atrophy and the findings of novel therapeutics to obviate this deleterious process.

We invented MyoRep technology ^33^. The model we created is based on reporter mouse technology which uniquely combines the presence of insulating sequences and a constitutively and ubiquitously open locus ^15,34^. MyoRep is able to sense early atrophy either *in vitro* and *in vivo*; moreover, it can sense atrophy initiated either by local denervation or systemic cancer. MyoRep mice are able to emit a local bioluminescent signal easily detectable by *in vivo* imaging upon muscle denervation already one day after cut of the sciatic nerve, when muscles were not reduced in size yet, as also showed by Wei Li and coworkers ^11^ in a rat model where a reporter gene was knocked into an intron of the *MuRF1* gene. On top of that, MyoRep mice emit a signal that we found originating from back muscles earlier than other muscles when injected a cachexia-promoting tumor as the sarcoma MCG101, indicating a preferential loss of back muscles over others. Instead, reporter cells locally injected reveal atrophy in nude mice bearing human pancreatic tumors, but only in a local way and with no precocity ^35^. Similarly, the reporter mouse useful to monitor proteasome activity detects a late and not early event in the atrophic process ^10^.

An important advantage of MyoRep mouse over existing reporter mice and systems is that it is insensitive to circadian rhythms. MyoRep mice are in fact insensitive to atrophy upon fasting or upon diurnal-nocturnal variations of muscle size. This was obtained by deleting the GRE sequence by MuRF1 promoter. The other existing reporter systems still hold GRE in their regulatory sequences, making them unable to discriminate between pathological from physiological atrophy. For example, the reporter mouse described above and generated by Wei Li and colleagues is highly responsive to dexamethasone injections ^11^. A common set of genes namely atrogenes drive the atrophic processes in either physiological and pathological atrophies, making challenging the possibility of distinguishing among them. We believe that a GRE-containing reporter mouse is not suitable for *in vivo* experiments aimed at identifying early biomarkers of atrophy or *ad hoc* drugs because more amenable of non-specific and undesirable activation (as we also showed by higher basal TWIST-based emissions in PBS-injected mice). Indeed, we provide data indicating that glucocorticoids are increased in plasma of mice either upon fasting or injected with a non-cachectic tumor (4T1 that we showed unable to cause atrophy and unable to increase plasma pro-inflammatory IL-6 cytokine ^26^), further supporting their involvement in physiological atrophy as well as during tumor progression with no cachexia.

As other reporter *in vivo* systems to sense atrophy, MyoRep retains some advantages as the possibility to spare animals during experiments because each animal can be followed over time in longitudinal studies without the need to sacrifice mice at various timepoints to measure muscle size and atrogene expression. In the past, we have successfully used microCT to follow *in vivo* atrophy initiated by C26 tumors or ALS ^36^, but that system was more time-consuming and labor-intensive and did not allow to visualize in advance atrophy before it was already present in mice. Nonetheless, both systems are in accordance with the 3R rule (Replacement, Refinement, Reduction) as recommended by ^37, 14, 38^.

### Future applications for the MyoRep technology

MyoRep tool better than others may serve to identify not only early biomarkers of atrophy in plasma and muscles but also to better understand the dynamics of muscle involved, in other words which muscle are more prone to different kinds of pathological atrophy by their emission in *in vivo* imaging. We plan to cross MyoRep mouse with other disease models as SOD1G93A mice for ALS or other models for motoneuron diseases or hereditary myopathies as Marinesco Sjögren syndrome. MyoRep could be also subjected to partial or total body paralysis for severe spinal cord injury or in the Intensive Care Unit (ICU) model ^39^, or to mechanical local muscular trauma. MyoRep could serve to unravel potential anti-atrophic effect of life-style changes, for example various dietary interventions and/or physical activity sessions of various aerobic or anaerobic exercise protocols.

The inability of MyoRep to sense circadian rhythms variations and its feature to hold transcription factor-binding sites conserved across species, including humans, make it suitable to generate clinical grade MyoRep-AAV9 for early diagnostics in human patients, as done for other therapeutic AAV9 ^40, 41^. We may consider treating MyoRep cells also with human fluids as plasma or interstitial cancerous fluids or in the future aerosol-derived samples from cancer patients. This would help to identify in advance patients at risk of cancer cachexia, for example, in order to intervene on time with proper countermeasures as electrical muscle stimulation, exercise or nutritional supplementations ^42^, by sparing economic resources towards only those who may receive some real advantages. We are about to screen *in vitro* plasma from cancer patients to see if MyoRep-expressing myoblasts may predict in advance if they will develop atrophy.

MyoRep can sense more severe atrophy of cancer-bearing males with respect to females at least in Apc^Min/+^ mice, helping us to understand sex-specific mechanisms of atrophy to design novel therapies suited for sex, as we have showed for musclin. We have also reported the reversibility of the MyoRep induction by means of musclin-AAV9 injections even if only locally administered. This indicates that such technology can be switched off by anti-atrophic molecules and MuRF1 promoter is at once a marker to follow atrophy, even in its early stages, and a way to see drug response and eventually in the future to find effective drug doses, as we did in preliminary experiments not shown to identify the right amount of AAV9 to administer.

Since *MuRF1* gene gets activated also in heart during its atrophy in the so-called cardiac cachexia process ^43^ and given that actin promoter of B6.Cg-Tg(ACTA1-cre)79Jme/J mice used to restrict the expression of MyoRep only in skeletal muscle is expressed also in heart, we could follow in future studies MyoRep expression in the heart following various diseases. Cardiac MyoRep induction should be easily visualized because localized to a restricted region in the thorax. Nonetheless, we may plan to restrict MyoRep expression only in the heart by crossing the mouse with the stop codon with a mouse expressing Cre recombinase under a specific promoter for cardiac expression only (as the B6.FVB-Tg (Myh6-cre) 2182Mds/J mouse). Their derived progeny may serve to understand mechanisms at the basis of heart failure obtained with coronary artery ligation or other microsurgeries in animal models ^44^.

### Limitations of the MyoRep technology

Despite having two reporter genes under the MyoRep promoter, *Firefly Luciferase* and *tdTomato*, we never visualized in optical microscopy *in vivo* neither in muscles dissected and analysed for their fluorescence *ex vivo* the expression of *tdTomato* for reasons that deserve further experiments. The second reporter gene is separated by the first one by P2A linker that perhaps allows too low expression of the second one (i.e., tdTomato) to be detected.

Another limitation of this tool is that *MuRF1* gene is not induced in all kinds of atrophy as that associated to microgravity in spaceflight ^45^ or in Duchenne Muscular Dystrophy (DMD) ^46^, making useless MyoRep to study these types of atrophy. Finally, the resolution of *in vivo* imaging is not enough to discriminate among different muscles, but only to identify grossly their position, and needs to be coupled with higher resolution imaging (microCT) or *ex vivo* analysis of separated muscles to better understand the origin of emitted signal in MyoRep mice.

On a different note, current MyoRep mouse is on C57BL/6J background that while offering the advantage to cross it with many disease models, has the disadvantage that black hair can mask bioluminescence, so that we shaved the mice before imaging them. This problem could be circumvented by generating albino MyoRep mice or by systemically injecting MyoRep-AAV9 in mice with white hair (i.e., Balb/C mice).

Despite these limitations, we believe that MyoRep technology constitutes a real advancement in the technology to study *in vivo* muscle wasting because able to discriminate successfully between pathological and physiological atrophy.

## Supporting information

6 Supplementary Figures with legends

## AVAILABILITY

Further details to the MyoRep technology can be found here: https://www.knowledge-share.eu/en/patents/reporter-system-for-muscle-atrophy https://patentscope.wipo.int/search/en/detail.jsf?docId=WO2022054012&_cid=P11-M73HPP-93383-1

## ACKNOWLEDGMENTS

We are grateful to Dr. Lorenza Ronfani and her team members from the Core Facility for Conditional Mutagenesis, San Raffaele Institute, Milan, Italy, for technical assistance in the generation of the MyoRep mouse.

