## Supplementary material for "MyoRep: a novel reporter system to detect early muscle atrophy *in vitro* and *in vivo*": 6 Supplementary Figures with legends

Supplementary Figure 1

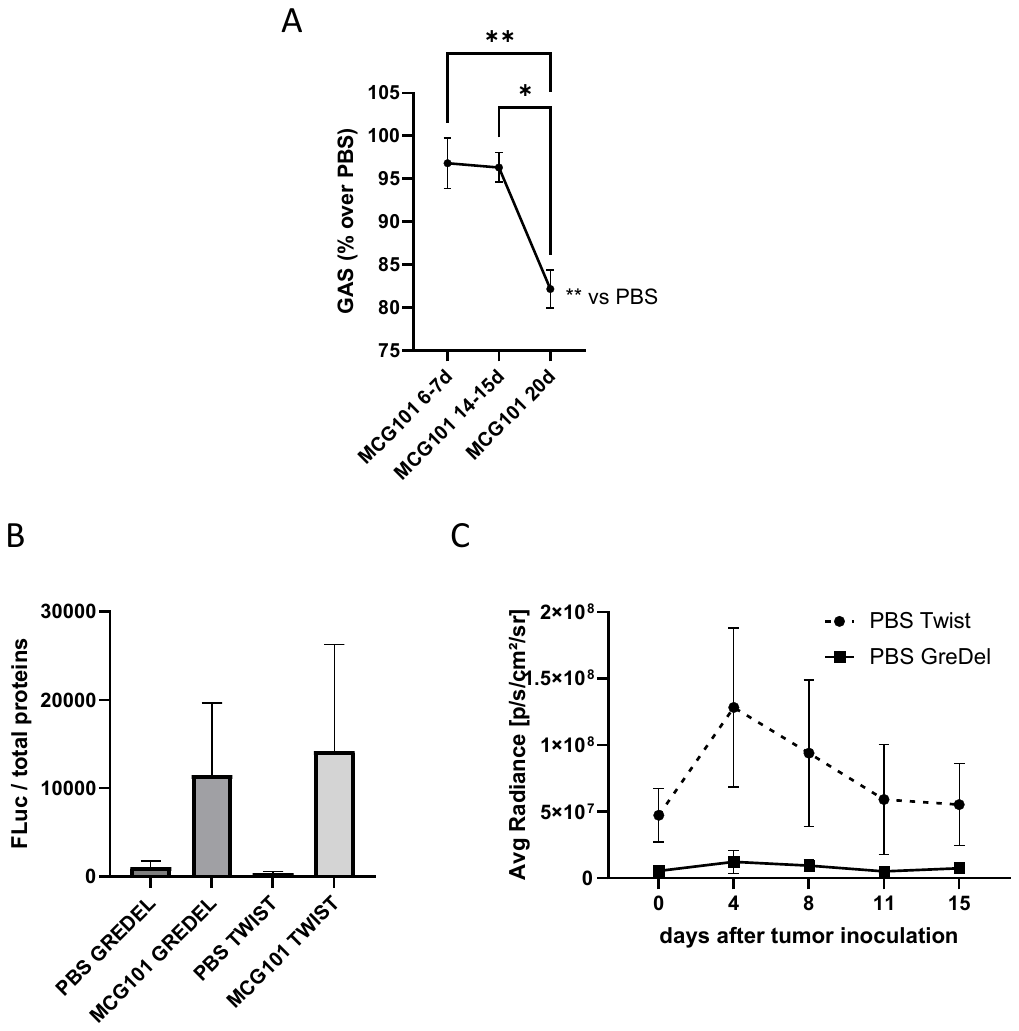

**Supplementary Figure 1** - (**A**) Gastrocnemius (GAS) weight (% over PBS) of MCG101 bearing-mice sacrificed at different times. Kruskal-Wallis’s test followed by Dunn’s post hoc test. *p≤0.05, **p≤0.01. MGC101 20d *vs* PBS, Unpaired t test, **p≤0.01. N = 7-33. (**B**) Luciferase Assay analysis of TA from mice of Figure 2. Total proteins were used to normalize the data. N = 4-5. All data are reported as mean ± SEM. (**C**) Analysis of photon emission comparing area under the curve (AUC) of TA expressing TWIST reporter *vs* TA expressing GREDEL reporter in PBS bearing-mice. Unpaired t test for AUC, **p≤0.01, N = 5.

Supplementary Figure 2

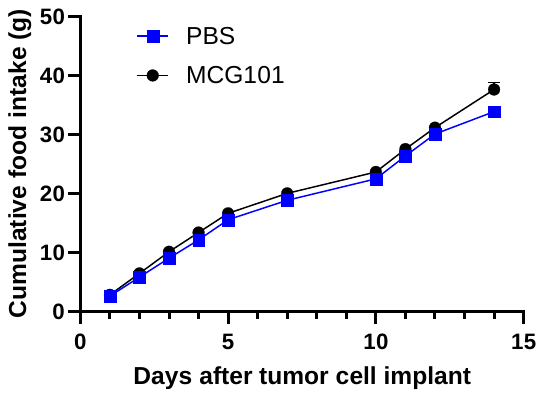

**Supplementary Figure 2 -** Cumulative food intake is shown for MCG101- and PBS-injected mice (cages = 1-2). Ns, multiple t test.

Supplementary Figure 3

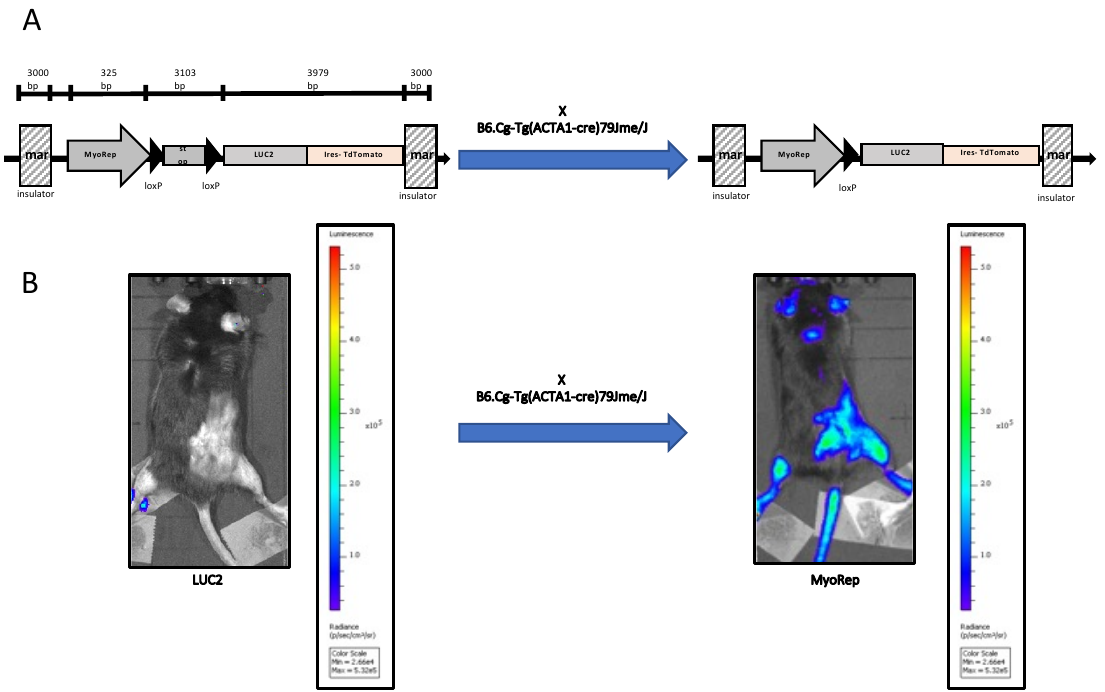

**Supplementary Figure 3 -** (**A**) A scheme for the generation of the MyoRep mouse. The stop sequence in Luc2 mouse is removed using the loxP system after crossing with the B6.Cg-Tg(ACTA1-cre)79Jme/J mouse. (**B**) Compared to the Luc2 mouse, the MyoRep mouse shows a basal bioluminescence signal, following the removal of the stop sequence.

Supplementary Figure 4

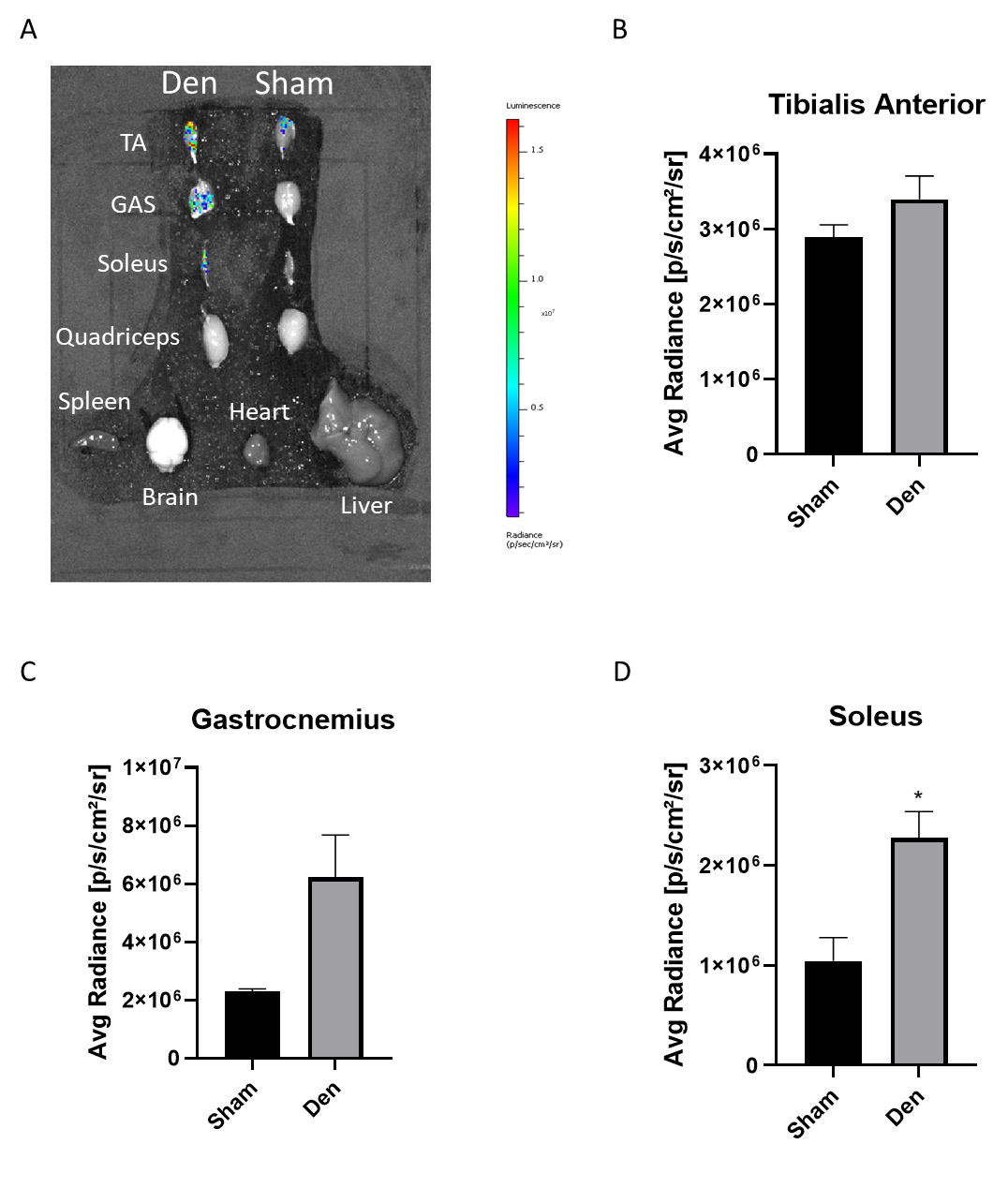

**Supplementary Figure 4 -** (**A**) *Ex vivo* imaging of muscles and organs from MyoRep mice subjected to denervation or sham operation. (**B-D**) Analysis of photon emission of the indicated muscles from MyoRep mice comparing denervated leg with sham-operated contralateral one. Mann Whitney test. *p≤0.05. N = 2. All data are reported as mean ± SEM.

Supplementary Figure 5

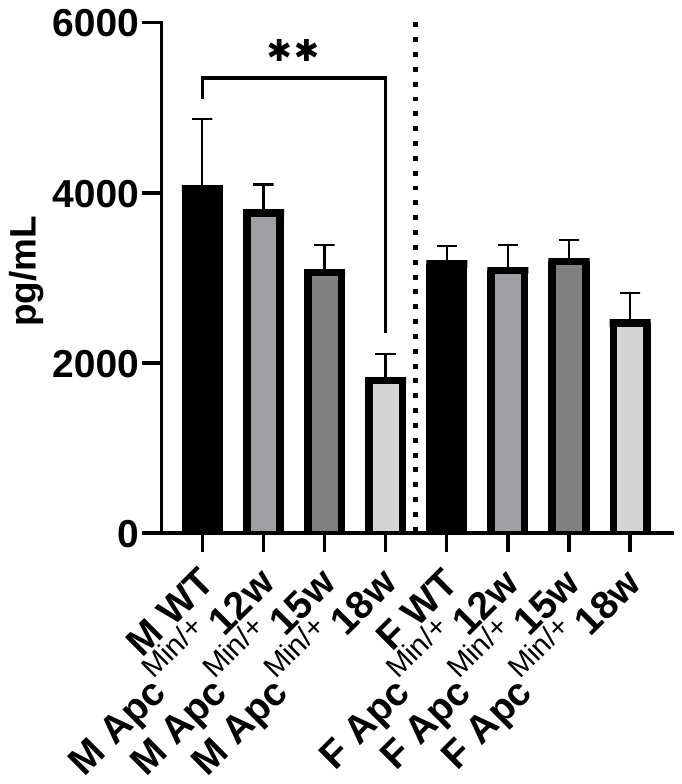

**Supplementary Figure 5 -** Levels of musclin have been measured through ELISA in plasma of WT and Apc^Min/+^ male and female mice at 12, 15, and 18 weeks of age. WT include samples of mice at the all indicated ages that have been pooled together. Kruskall-Wallis test followed by Dunn’s post-hoc test, **p ≤0.01. N =10. All data are reported as mean ± SEM.

Supplementary Figure 6

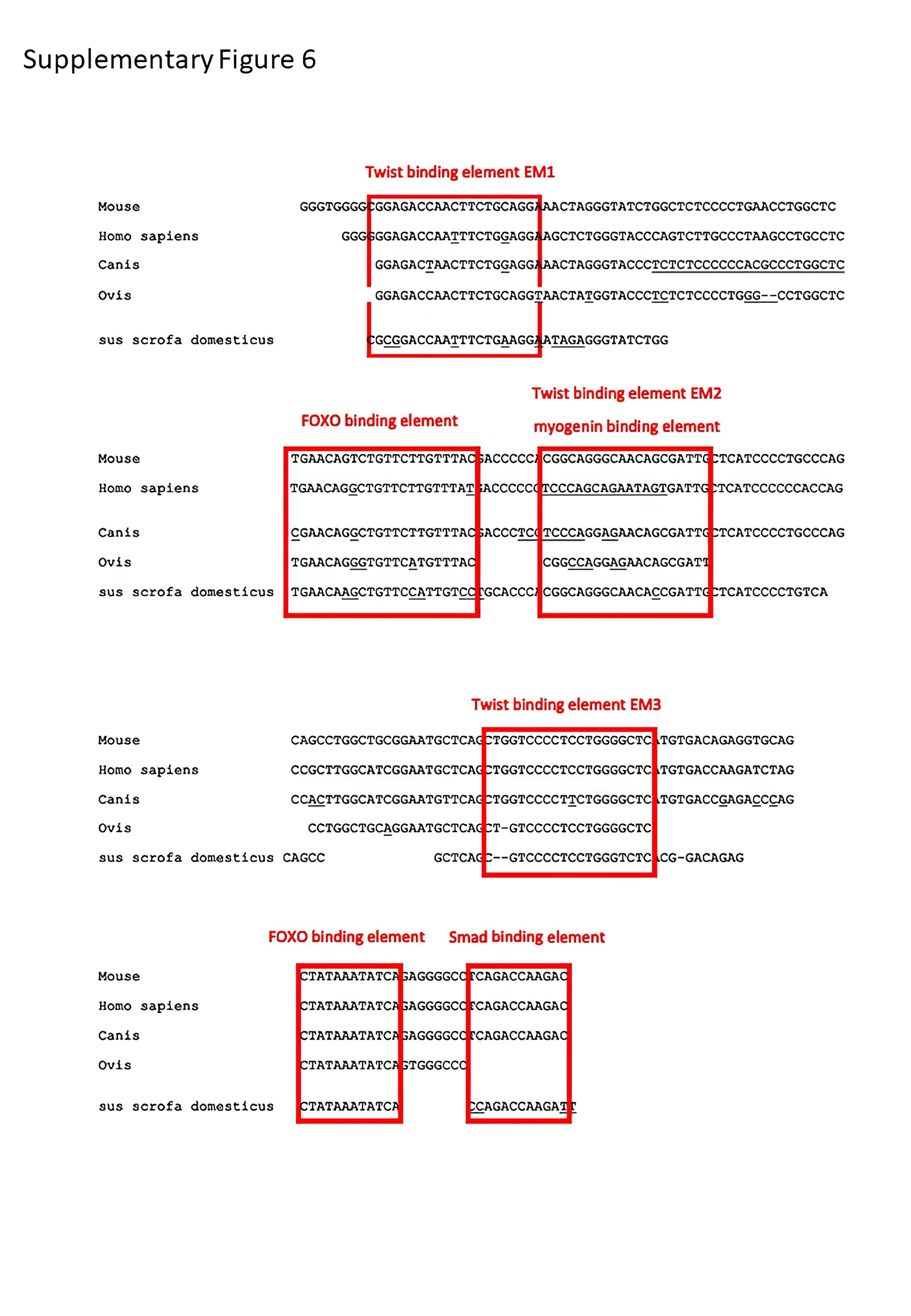

**Supplementary Figure 6 -** The alignment of transcription factor-binding sites, boxed in red, conserved across species are shown in the MyoRep promoter sequence. Analysis done with BLAST database.
